# State-Based Delay Representation and Its Transfer from a Game of Pong to Reaching and Tracking

**DOI:** 10.1101/195982

**Authors:** Guy Avraham, Raz Leib, Assaf Pressman, Lucia S. Simo, Amir Karniel, Lior Shmuelof, Ferdinando A. Mussa-Ivaldi, Ilana Nisky

## Abstract

To accurately estimate the state of the body, the nervous system needs to account for delays between signals from different sensory modalities. To investigate how such delays may be represented in the sensorimotor system, we asked human participants to play a virtual pong game in which the movement of the virtual paddle was delayed with respect to their hand movement. We tested the representation of this new mapping between the hand and the delayed paddle by examining transfer of adaptation to blind reaching and blind tracking tasks. These blind tasks enabled to capture the representation in feedforward mechanisms of movement control. A Time Representation of the delay is an estimation of the actual time lag between hand and paddle movements. A State Representation is a representation of delay using current state variables: the distance between the paddle and the ball originating from the delay may be considered as a spatial shift; the low sensitivity in the response of the paddle may be interpreted as a minifying gain; and the lag may be attributed to a mechanical resistance that influences paddle’s movement. We found that the effects of prolonged exposure to the delayed feedback transferred to blind reaching and tracking tasks and caused participants to exhibit hypermetric movements. These results, together with simulations of our representation models, suggest that delay is not represented based on time, but rather as a spatial gain change in visuomotor mapping.

**Significance Statement:** It is known that the brain copes with sensory feedback delays to control movements, but it is unclear whether it does so using a representation of the actual time lag. We addressed this question by exposing participants to a visuomotor delay during a dynamic game of pong. Following the game, participants exhibited hypermetric reaching and tracking movements that indicate that delay is represented as a visuomotor gain rather than as a temporal shift.

## Introduction

It is unclear if the brain represents time explicitly (Karniel, 2011) using “neural clocks” (Ivry, 1996; Spencer et al., 2003; Ivry and Schlerf, 2008). Evidence suggests that no such clock is involved in the control of movement: humans can adapt to force perturbations that depend on the state of the arm (position, velocity, etc.), but not to forces that are explicit functions of time (Karniel and Mussa-Ivaldi, 2003); also, time-dependent forces are sometimes treated as state-dependent (Conditt and Mussa-Ivaldi, 1999). Instead, for the timing of movements, the sensorimotor system may use the temporal dynamics of state variables that are associated with the performance of actions.

Time representation is important for sensory integration, movement planning and execution. Sensory signals are characterized by different transmission delays (Murray and Wallace, 2011), and movement planning and execution require additional processing time. Therefore, to enable the organism’s survival, the sensorimotor system must account for these delays. The current literature is equivocal on how delays are represented. Humans can adapt to visuomotor delays (Miall and Jackson, 2006; Botzer and Karniel, 2013) and to delayed force feedback (Witney et al., 1999; Levy et al., 2010; Leib et al., 2015; Avraham et al., 2017). However, delayed feedback biases perception of impedance (Pressman et al., 2007; Nisky et al., 2008; Nisky et al., 2010; Di Luca et al., 2011; Kuling et al., 2015; Takamuku and Gomi, 2015; Leib et al., 2016), suggesting that the sensorimotor system has limited capability to realign the signals for accurate estimations of the environment (Ionta et al., 2014).

To understand how the sensorimotor system controls movements with non-synchronized feedback, we examined the representation of visuomotor delay in an ecological interception task. Participants played a pong game and controlled a paddle to hit a moving ball. The paddle movement was either coincident or delayed with respect to hand movement (Fig. 1). Because the delay influences the distance between the hand and the paddle, its representation can be *Time-based* or *State-based*. In *Time Representation*, the player represents the actual time lag whereas in *State Representation*, she uses current state variables, and may attribute the distance between the hand and the paddle to a spatial shift, a minifying gain, or a mechanical resistance. Using *Time Representation*, the player would precede the movement of the hand by the appropriate time so that the paddle would hit the ball at the planned location. Instead, using *State Representation*, she would aim her hand to a farther location.

**Fig 1.**
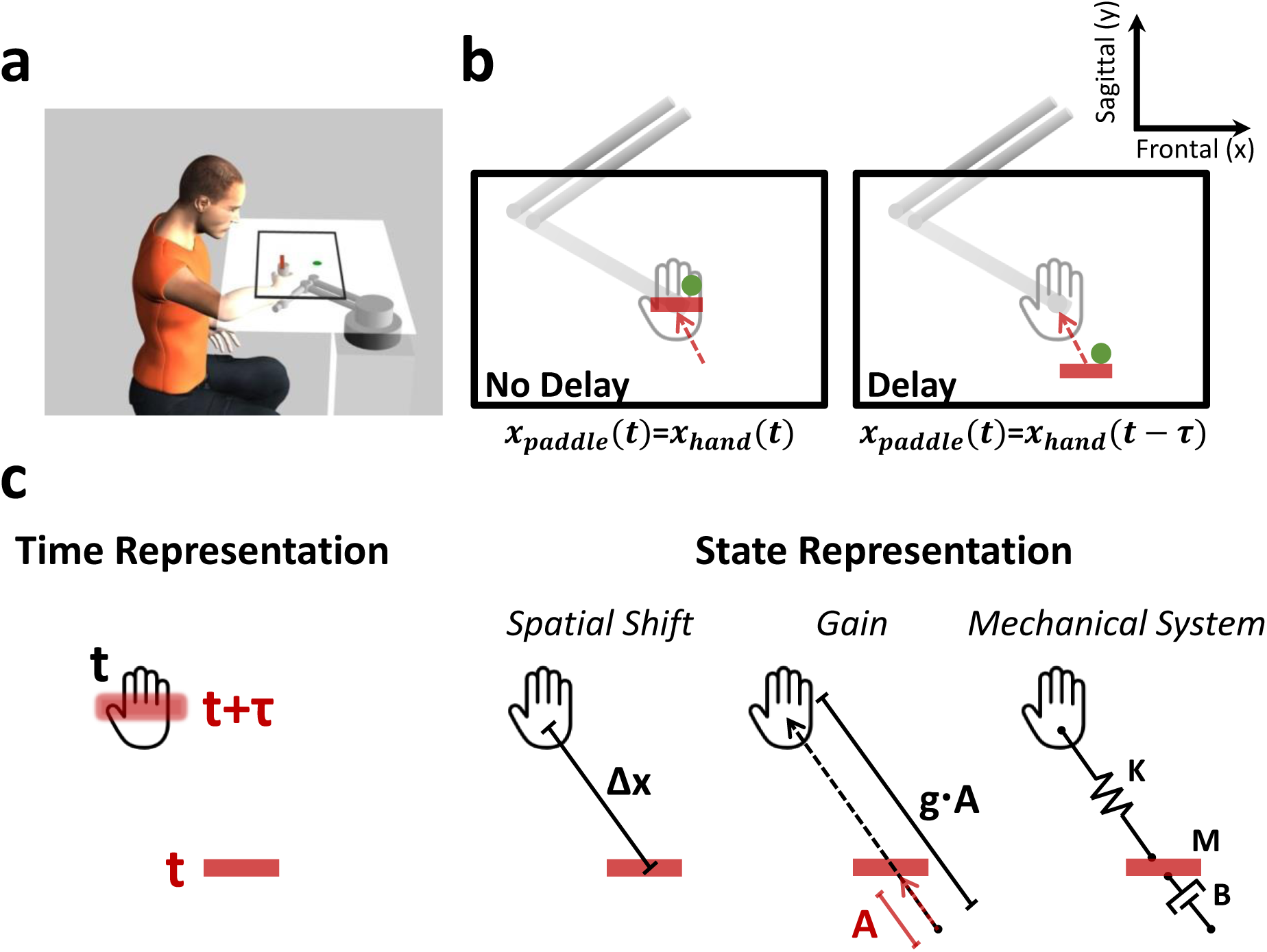
The pong game and the representation models for hand-paddle delay. (a) An illustration of the experimental setup and the pong game: participants sat and held the handle of a robotic arm. A screen that was placed horizontally above their hand covered the hand and displayed the scene of the experiment. During the pong game, participants controlled the movement of the paddle (red bar) and were required to hit a moving ball (green dot) towards the upper wall of the pong arena, which is delineated by the black rectangle. (b) The paddle movement was either concurrent (left – No Delay) or delayed (right – Delay) with respect to the hand movement (the red arrow indicates the paddle movement direction). (c) Participants could represent the hand location based on the delayed paddle using a *Time Representation* (left) or a *State Representation* (right). In a *Time Representation*, participants were assumed to estimate the actual time lag, *Τ*, and represented the hand location at time *t* as the location of the paddle at *t* + *Τ?*(blurred paddle). In a *State Representation*, participants would represent a *Spatial Shift* (Δ*x*) between the hand and the paddle, an altered visuomotor *Gain* (*g*) relationship between hand and paddle movements, or a *Mechanical System* that connects the two and includes a spring (**K**), a mass (**M**) and a damper (**B**).

Coping with delayed feedback is critical for forming internal representations in feedforward control. A thorough understanding of this process requires identifying delay effects on feedforward mechanisms of movement coordination. Such mechanisms can be isolated only in the absence of visual feedback. Previous studies suggested the *Time-based* (Rohde et al., 2014; Farshchiansadegh et al., 2015) and the *State-based* spatial shift (Smith and Bowen, 1980) and mechanical system (Sarlegna et al., 2010; Takamuku and Gomi, 2015; Leib et al., 2017) as candidate representation models for visuomotor delay (Rohde and Ernst, 2016). They probed delay effects on perception, or by observing action during adaptation and its aftereffects, but always with visual feedback. Also, these studies evaluated the representation using a single task. Another common approach to characterizing changes in internal representations is to examine transfer of adaptation (Shadmehr and Mussa-Ivaldi, 1994; Krakauer et al., 2006). While various terminologies are used in different fields, we define transfer as a change in performance in one task after experiencing another task. We explored visuomotor delay representation in the pong game by investigating its transfer to blind reaching and tracking tasks. Transfer to these well-understood movements allowed for comparing our experimental observations to simulations of the four representation models. By omitting the visual feedback, we could examine performance when participants had to rely solely on feedforward control and proprioceptive feedback. Thus, our transfer tasks enabled capturing visuomotor delay representation in feedforward mechanisms.

Abrupt or gradual perturbation schedule was shown to affect transfer of adaptation. Specifically, stronger transfers were reported after gradual presentations (Kluzik et al., 2008; Torres-Oviedo and Bastian, 2012), possibly due to the influences of awareness (Kluzik et al., 2008) and credit assignment (Berniker and Kording, 2008). We hypothesized that a gradual rather than an abrupt increase in the delay during the game would enhance the behavioral effects in our transfer tasks.

Our simulations and experimental results suggest a state-based visuomotor delay representation that is not influenced by perturbation schedule. Particularly, performance changes in both pong and transfer tasks favor a delay representation as a gain change in visuomotor mapping.

## Methods

### Notations

We use lower-case letters for scalars, lower-case bold letters for vectors, and upper-case bold letters for matrices. **x** is the Cartesian space position vector, with *x* and *y* the position coordinates (for the right-left / frontal and forward-backward / sagittal planes, respectively). **F** is the force vector, with *f _x_* and *f _y_* force coordinates. N indicates the number of participants in a group.

### Experiments

#### Participants and experimental setup

Seventy-seven healthy volunteers (aged [19-41], 41 females) participated in four experiments: 17 participated in Experiment 1, 20 in Experiment 2, 20 in Experiment 3 and 20 in Experiment 4. Human subjects were recruited at a location which will be identified if the article is published.

The experiments were administered in a virtual reality environment in which the participants controlled the handle of a robotic device, either a six degrees-of-freedom PHANTOM^®^ Premium^™^1.5 haptic device (Geomagic^®^) (Experiments 1 and 4), a two degrees-of-freedom MIT Manipulandum (Experiment 2) or a six degrees-of-freedom PHANTOM^®^ Premium^™^ 3.0 haptic device (Geomagic^®^) (Experiment 3). Figure 1a illustrates the experimental setup. Seated participants held the handle of the device with their right hand while looking at a screen that was placed horizontally above their hand, at a distance of ∼10 cm below the participants’ chin. They were instructed to move in a horizontal (transverse) plane. In Experiments 1, 3 and 4, hand position was maintained in this plane by forces generated by the device that resisted any vertical movement. The update rate of the control loop was 1,000 Hz. Since the Manipulandum is planar, this was not required in Experiment 2. In Experiments 1, 2 and 4, a projector that was suspended from the ceiling projected the scene onto a horizontal white screen placed above the participant’s arm. In Experiment 3, a flat LED television was suspended approximately 20 cm above a reflective screen, placing the visual scene approximately 20 cm below the screen, on the horizontal plane in which the hand was moving. The hand was hidden from sight by the screen, and a dark sheet covered the upper body of the participants to remove all visual cues about the arm configuration. When visual feedback of the hand location was provided, the movement of the device was mapped to the movement of a cursor; when it was not perturbed by the delay, the cursor movement was consistent with the hand movement, with a delay of 5 (Experiment 2) or 10 (Experiments 1, 3 and 4) ms due to the refresh rate of the display. The experimentally manipulated delay in the delay condition was added on top of this delay.

#### Tasks

Each experiment consisted of two tasks: a pong game task and another “blind” task. During the latter, no visual feedback about the hand location was provided. In Experiments 1 and 2, the blind task was a reaching task, and in Experiments 3 and 4, it was a tracking task. The purposes of the blind tasks were to examine transfer and to capture the participants’ representation of the hand-cursor dynamics following exposure to either the non-delayed or delayed pong game.

##### Pong game

In the pong game, participants observed the scene illustrated in Figure 1b. The rectangle delineated by the black walls (Experiments 1 and 3: [sagittal × frontal dimensions] 16 × 24 cm, Experiment 2: 17 × 34 cm, Experiment 4: 18 × 26 cm) indicates the pong arena. The red horizontal bar marks the location of the paddle and corresponds to the hand location. As described below (see **Protocol**), each experiment consisted of two Pong game sessions. We termed the first Pong session **Pong No Delay**, and the second Pong session **Pong Delay**. In the **Pong No Delay** session, the paddle moved synchronously with the hand. In the **Pong Delay** session, the paddle movement was delayed with respect to the hand movement: **x** _*p*_ (*t*) = **x**_*h*_ (*t* −*Τ*), where **x** _*p*_ (*t*) and **x**_*h*_ (*t*) are the positions of the paddle and the hand, respectively, and *Τ* is the applied delay (note that for the Control group in Experiment 1 alone, the delay in the **Pong Delay** session was equal to zero, and hence, the dynamics between the hand and the paddle in this session was equivalent to the dynamics during the **Pong No Delay** session). To apply the delay, we saved the location of the hand in a buffer that was updated with the update rate of the control loop, and displayed the paddle at the location of the hand *Τ* time prior to it. *Τ* was set to values between 0 and 0.1 s, depending on the protocol and the stage within the session. The green dot indicates a ball that bounces off the walls and the paddle as it hits them. The duration of each Pong trial was *t*_*Trial*_ = 60 *s*. Information about the elapsed time from the beginning of the trial was provided to the participants by a magenta-colored timer bar. Feedback on performance in each trial was also provided using a blue hit bar that incremented according to the recorded paddle-ball hits from trial initiation onward. In Experiments 1, 3 and 4, during each trial, we updated the hit bar on every hit. The total amount of hits required to fill the bar completely 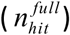 was set to 80 in Experiment 1 and to 60 in Experiments 3 and 4, and it remained constant throughout the entire experiment. In Experiment 2, during each trial, we updated the hit bar every time the participants reached 5% of 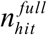 During the **Pong No Delay** session, we set 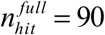. After the last trial of the **Pong No Delay** session, we calculated each participant’s average hitting rate on that trial, *n*_*hit*_/*t*_*Trial*_ where *n*_*hit*_ is the number of hits on the last trial of the **Pong No Delay** session. On the first trial of the second **Pong Delay** session, we matched the progression rate of the hit bar for each participant according to performance at the end of the **Pong No Delay** session, such that 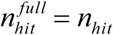 Then, in order to encourage participants to improve, we decreased the progression rate of the hit bar by 5% on each successive trial.

The ball was not displayed between trials. The initiation of a trial was associated with the appearance of the ball in the arena. In Experiments 1 and 2, a trial was initiated when participants moved the paddle to the “restart zone” – a green rectangle (Experiment 1: 1 × 4 cm, Experiment 2: 2 × 10 cm) that was placed 3 cm below the bottom (proximal) border of the arena. Throughout the entire experiment, including the **Pong Delay** session, the paddle was never delayed between trials. Since the displayed paddle movement between trials was always instantaneous with hand movement, we were concerned that the effect of delay on state representation could be attenuated by a recalibration of the estimated hand location according to the non-delayed paddle. Thus, in Experiments 3 and 4, we did not display the paddle between trials, and participants were instructed to initiate a trial by moving the handle of the robotic device backward (towards their body). When the invisible paddle crossed a distance of 3 cm from the bottom border of the arena, the trial was initiated. In Experiments 1, 3 and 4, the initial velocity of the ball in the first Pong trial was 20 cm/s, and in every other Pong trial, it was the same as the velocity at the end of the previous trial. In Experiment 2, the initial velocity of the ball in each Pong trial was 28 cm/s.

The participants were instructed to hit the ball towards the upper (distal) wall as many times as possible. When the ball hit a wall, its movement direction was changed to the reflected arrival direction, keeping the same absolute velocity (consistent with the laws of elastic collision). To encourage the participants to explore the whole arena and to eliminate a drift to stationary strategies, the reflection of the upper wall (and not the other walls) included some random jitter. Introducing the jitter effectively corresponded to creating a compromise between playing against a wall and playing against an opponent. This was done by adding the jitter component *j* to the horizontal component of the ball velocity before the collision with the upper wall 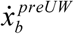, such that:

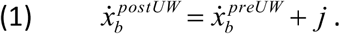

where 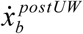 is the horizontal component of ball’s velocity following the collision with the upper wall. In Experiments 1, 3 and 4, 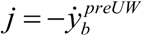 tan(α_*j*_), where 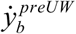 is the vertical component of ball’s velocity before the collision with the upper wall and *α* ∼ *N*(μ,*σ* 2)= *N*(0,0.05*π*) *μ* and *σ*2 denote the mean and variance of the normal distribution *N*, respectively. In Experiment 2, 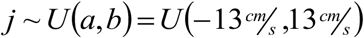, where *U* is the uniform distribution between its two arguments.

The velocity of the ball was also influenced by the paddle velocity at the time of a hit. We determined the relationship between the velocity of the ball following a paddle hit 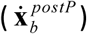 according to the velocity of the ball before the hit 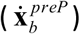 and the velocity of the paddle when contacting the ball 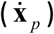 For the frontal dimension, the ball velocity after bouncing off the paddle was computed as:

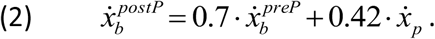

For the sagittal dimension, we let a hit occur only when the paddle was moving upward and the ball was moving downward. In all other cases, the ball passed through the paddle as if it was moving over different planes. The rationale for allowing hits to occur only in the upward direction was to differentiate between the effects of the *Time* and *State – Spatial Shift* representation models. In our design, we assumed that a change in representation would occur primarily during meaningful events in the pong game; i.e., paddle-ball hits. Hence, allowing hits to occur in both the upward and downward directions could have cancelled the *State Representation – Spatial Shift* effect, and would have restricted our ability to distinguish it from the *Time Representation* model. In this dimension, after a hit occurred, the ball’s movement direction was always reversed, and its velocity was computed as:

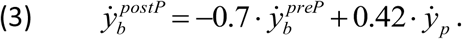

In our setup, the forward movement direction had a positive velocity, and the backward direction was negative. Note that since a hit occurred only when 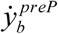 was negative and 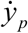 was positive, the resulting 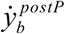 was always positive. This way, the ball movement direction following the hit was reversed, and moved towards the upper wall. A possible strategy to cope with the delay was to slow down, and thus, for the delay to be effective, we encouraged participants to maintain their movement velocity as much as possible during the game despite the change in delay. Therefore, we set the coefficients’ absolute values of 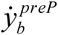 and 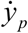 (Eq. 3) to be between 0 and 1, such that they would reduce the effect of these velocities on the velocity of the ball after the hit. Thus, to maintain the ball speed after the hit as it was before the hit or to make it faster, 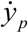 needed to be at least 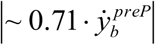 In addition to the constraint on the paddle to move upward, participants were informed that they should control the paddle to move fast enough at the moment of a hit, otherwise the ball would slow down, reducing the number of opportunities to hit it.

Once participants hit the ball with the paddle, a haptic pulse was delivered by the device simultaneously with the displayed collision; that is, when the paddle was delayed, the pulse was delayed. This design was thought to strengthen the delay effect during the hit. The pulse **f** ^*postP*^ was applied according to:

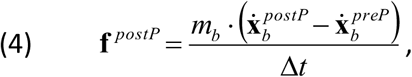

where *m*_*b*_ is the ball’s mass and Δ*t* is the duration of the applied force. The specific parameters of the magnitude and durations of the haptic pulses were tuned for each of the devices that were used in the different experiments such that a relatively similar haptic stimulation was applied despite the differences in the specifications of the devices. In Experiments 1, 3 and 4, *m b* = 0.15 *kg*, and we calculated **f** ^*postP*^ as the maximum applied force according to a time interval of Δ*t* = 0.025 *s*. However, the haptic pulse was applied for 0.05 *s*, in which it gradually and linearly increased from zero to **f** ^*postP*^ for the first 0.025 *s* (since the update rate of the control loop in this setup was 1,000 Hz, this is equivalent to 25 sample intervals) and then decreased back to zero in a similar manner for the remaining 0.025 *s*. In Experiment 2, *m*_*b*_ = 0.05 *kg*, and the force was applied during a single sample interval of Δ*t* = 0.005 *s*.

##### Reaching

At the beginning of a reaching trial, the entire display was turned off, and the device applied a spring-like force that brought the hand to a start location, which was at the center of the bottom wall of the pong arena (that was displayed only during the pong trials) and 1 cm (Experiment 1) or 3 cm (Experiment 2) below it. A trial began when a target (a hollow square, 1.5 × 1.5 cm inner area) appeared in one of three locations in the plane, which were 10 cm (Experiment 1) or 12 cm (Experiment 2) from the start location in the forward direction, and separated from each other by 45^0^ (Fig. 2a,b, 4a and 6a). Throughout a reaching session, each of the three targets appeared fifteen times in a random and predetermined order. The appearance of the target was the cue for the participants to reach fast and to stop at the target. During each reaching trial in the experiment, we defined movement initiation as the time when the hand was 3 cm from the start location (Experiment 1) or when the sagittal component of the hand velocity 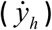 exceeded 25 cm/s (Experiment 2). Movement stop was defined at 0.5 s after 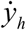 went below 10 cm/s (Experiment 1) or 0.2 s after it went below 15 cm/s (Experiment 2). After identifying that a reaching movement had been initiated and completed, the device returned the hand to the start location in preparation for the next target to appear. We used three types of reaching sessions that differed from each other in terms of the visual feedback provided to the participants (Fig. 2a and 2b). During the **Reach – Training** session, participants received full visual feedback on the hand location using a cursor (filled square, 1.5 × 1.5 cm) on the screen throughout the entire movement. They were instructed to put the cursor inside the hollow target. During the **Blind Reach – Training** session, the cursor was not presented during the movement, and participants were requested to imagine there was a cursor, and to stop when the invisible cursor was within the target. When they stopped, we displayed the cursor, providing the participants with feedback about their movement endpoint with respect to the location of the target. During the **Blind Reach** sessions that were presented after each of the **Pong** sessions, participants did not receive any visual feedback about their performance during or after the trial.

**Fig 2.**
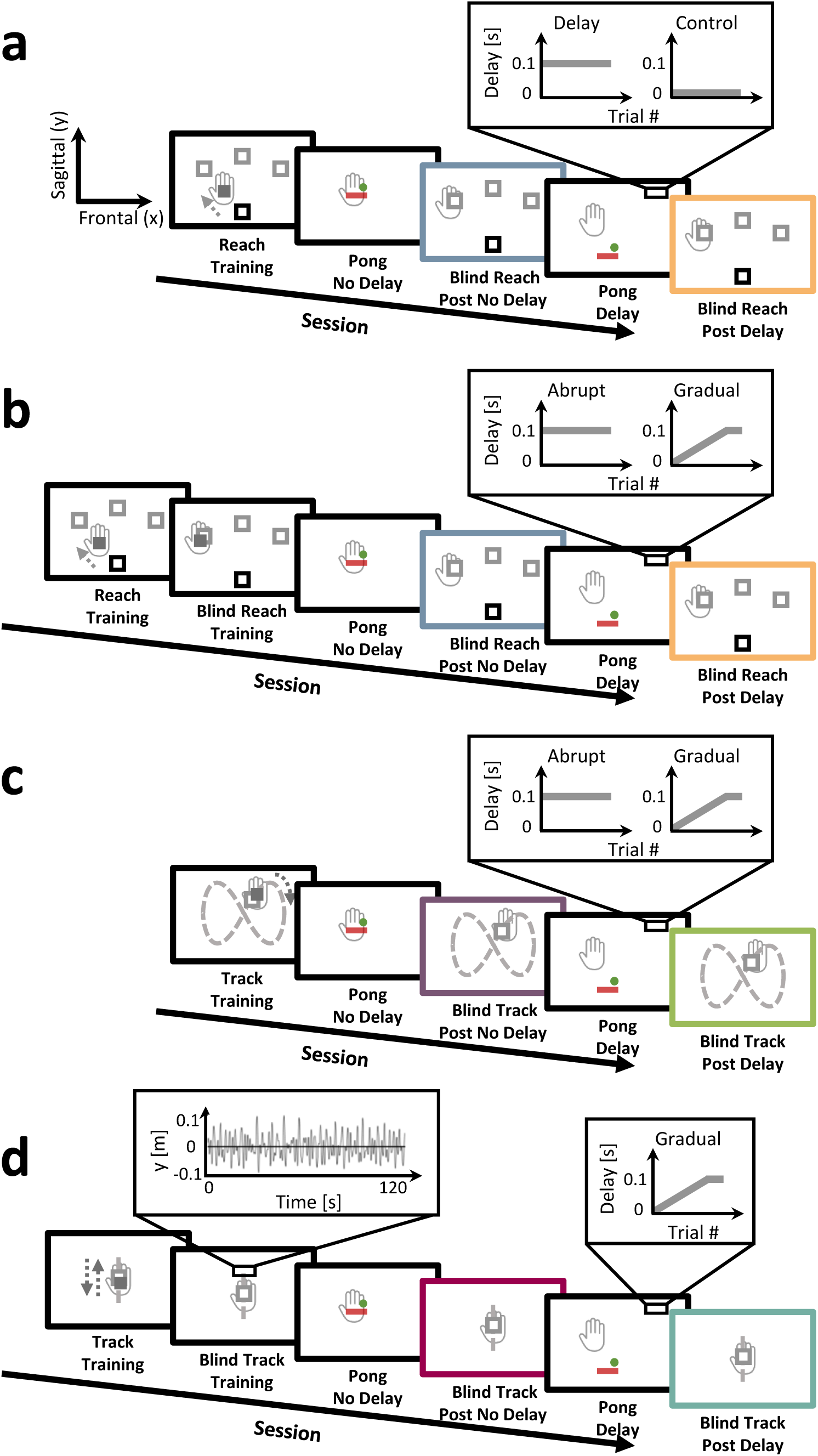
Experimental protocols.

##### Tracking – figure-of-eight

At the beginning of a tracking trial, the entire display was turned off, and the device applied a spring-like force that brought the hand to a start location, which was at the center of the bottom wall of the Pong arena and 2 cm below it. During each trial, participants were asked to track a target (a hollow square, 1.5 × 1.5 cm inner area) that moved along an invisible figure-of-eight path (Fig. 2c). This path was constructed as a combination of the following cyclic trajectories in the two-dimensional plane:

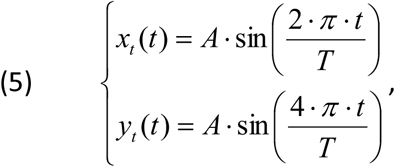

Where *A* = 8 *cm* is the path amplitude, and *T* = 5 *s* is the cycle time. The center of the figure-of-eight path was located 15 cm ahead of the location of the hand at trial initiation (start location). A trial began when a target appeared in one of five locations in the plane: either in the center of the figure-of-eight path (15 cm ahead of the start location), or in each of the four sagittal extrema (∼9 and ∼24 cm ahead). Throughout a tracking session, the five targets appeared in equally often and in a random and predetermined order. The appearance of the target was the cue for the participants to reach fast and stop at the target. Reaching initiation was defined as the time when either the frontal 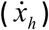 or sagittal 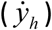 components of hand velocity exceeded 10 cm/s. Reaching stop was defined as 0.5 s after both 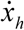 and 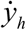 went below 5 cm/s. When the reaching movement stopped, the target started moving along the figure-of-eight path until it returned to its initial location. The targets moved in the same direction along the path (as illustrated by the dotted arrow in Fig. 2c), regardless of their initial location. A trial was completed after the device returned the hand to the start location in preparation for the next target to appear. Each experiment included two types of tracking sessions that differed from each other by the visual feedback that was provided to the participants (Fig. 2c). During the **Track – Training** session, participants received full visual feedback on the hand location using a cursor (filled square, 1.5 × 1.5 cm) on the screen throughout the entire movement. They were instructed to keep the cursor inside the hollow target. During the **Blind Track** session, the cursor was not visible during the trial, and participants were requested to imagine there was a cursor, and to keep the imagined cursor within the moving target.

##### Tracking – mixture of sinusoids

At the beginning of the tracking trial, the entire display was turned off, and the device applied a spring-like force that brought the hand to a start location, which was 2 cm above the center of the Pong arena. This was followed by the appearance of a target (a hollow square, 1.5 × 1.5 cm inner area) above the start location. A trial began two seconds later with the movement initiation of the target along an invisible one-dimensional path (Fig. 2d). This path was constructed as a mixture of five cyclic trajectories, all of which had the same amplitude (*A* = 2 *cm*) but each trajectory consisted of a different frequency (*fr* = [0.31, 0.67, 0.23, 0.42, 0.54]) and phase (*φ* = [0,*π* 4,*π*, 3*π* 2,*π* 3]):

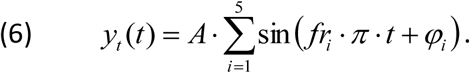

In each trial, participants were asked to track the movement of the target. The duration of each trial was two minutes. The tracking path was the same across trials. Each experiment included two types of tracking sessions that were different from each other in terms of the visual feedback that was provided to the participants (Fig. 2d). During a **Track – Training** session, participants received full visual feedback of their hand location using a cursor (filled square, 1.5 × 1.5 cm) on the screen throughout the entire movement. They were instructed to keep the cursor inside the hollow target. During the **Blind Track** sessions, the cursor was not presented, and participants were requested to imagine there was a cursor, and to keep the imagined cursor within the moving target.

#### Protocol

##### Experiment 1

In each experiment, sessions alternated a pong game and a reaching task (Fig. 2a). Each **Reach** session consisted of 45 trials (fifteen for each target). An experiment started with a **Reach – Training** session. The purpose of this session was to familiarize participants with the reaching task. After training, participants were presented with the **Pong No Delay** session for ∼10 min. This was followed by a **Blind Reach** session (**Post No Delay**). Next, participants experienced the **Pong Delay** session for ∼30 min. In the Delay group (N=9), we introduced a delay of *Τ* = 0.1 *s* between hand and paddle movements on the first trial of the **Pong Delay** session, which remained constant throughout the entire session. In the Control group (N=8), no delay was applied in **Pong Delay** session. The **Pong Delay** session was followed by another **Blind Reach** session (**Post Delay**).

In all experiments, the participants’ hand (gray) was hidden from sight the entire time. (a) Experiment 1: Delay vs. Control, transfer to reaching. Sessions alternated between a pong game and a reaching task. During a reach trial, a target (gray square) appeared in one of three locations in space beyond a start location (black square), and participants were asked to reach and stop at the target. An experiment started with a **Reach – Training** session in which participants received full visual feedback of the hand location using a cursor on the screen (dark gray filled square). After training, participants were presented with a **Pong** game session (**No Delay**), in which the paddle moved instantaneously with their hand movement, followed by a **Blind Reach** session where no visual feedback was provided at any point during the trial (**Post No Delay**, blue frame). The second **Pong** game session (**Delay**) was introduced with a delay (Delay group) or without a delay (Control group) between hand and paddle movements, and was followed by another **Blind Reach** session (**Post Delay**, orange frame). (b) Experiment 2: Abrupt vs. Gradual delay, transfer to reaching. The experimental protocol was similar to Experiment 1, but with the addition of a **Blind Reach – Training** session: the cursor was omitted during movement, but was displayed at the movement stop location. In the second **Pong** game session, we introduced either an abruptly (Abrupt group) or gradually (Gradual group) increasing delay. (c) Experiment 3: Abrupt vs. Gradual delay, transfer to tracking (figure-of-eight). Sessions alternated between a pong game and a tracking task. During a track trial, participants were asked to track a target that moved along a figure-of-eight path (dashed gray. The path was not presented to the participants) in a direction illustrated by the dotted dark gray arrow. The experiment started with a **Track – Training** session in which participants received full visual feedback on their hand location (dark gray filled square). After training, participants were presented with a **Pong** game session with no delay (**No Delay**), followed by a **Blind Track** session (**Post No Delay**, purple frame). Next, a **Pong** game session was introduced with either an abruptly (Abrupt group) or gradually (Gradual group) increasing delay (**Delay**), and was followed by another **Blind Track** session (**Post Delay**, green frame). (d) Experiment 4: Gradual delay, transfer to tracking (mixture of sinusoids). Sessions alternated between a pong game and a tracking task. During a track trial, participants were asked to track a target that moved along a sagittal path (dashed gray. The path was not presented to the participants). The target trajectory (left zooming window) was designed as a mixture of five sinusoids of different frequencies and phases. The experiment started with a **Track – Training** session in which participants received full visual feedback on their hand location (dark gray filled square), followed by a **Blind Track – Training** session. After training, participants were presented with a **Pong** game session with no delay (**No Delay**), followed by a **Blind Track** session (**Post No Delay**, magenta frame). Next, a **Pong** game session was introduced with a gradually increasing delay (**Delay**), and was followed by another **Blind Track** session (**Post Delay**, cyan frame).

##### Experiment 2

In each experiment, sessions alternated a pong game and a reaching task (Fig. 2b). An experiment started with a **Reach – Training** session that consisted of six trials (two for each target) and familiarized participants with the reaching task. The next session was a **Blind Reach – Training** session that consisted of 45 trials (15 for each target). By providing visual feedback only after the movement ended, we aimed in this session to train participants to reach accurately to the targets when they did not have any visual indication of their hand location throughout the movement and to improve their baseline performance. After training, participants were presented with a **Pong No Delay** session consisting of 10 trials. This was followed by a **Blind Reach** session (**Post No Delay**) with 45 trials. Next, participants experienced a **Pong Delay** session consisting of 30 trials. In the Abrupt group (N=10), we introduced a delay of *Τ* = 0.1 *s* between hand and paddle movements on the first trial of the **Pong Delay** session that remained constant throughout the entire session. In the Gradual group (N=10), we introduced a delay of *Τ* = 0.004 *s* on the first trial of the **Pong Delay** session and gradually increased it by 0.004 *s* on every trial until the 25^th^ trial of the session, when it reached to *Τ* = 0.1 *s*; then, the delay was kept constant for the remaining five trials in the session. The experiment ended with another **Blind Reach** session (**Post Delay**) of 45 trials.

##### Experiment 3

In each experiment, sessions alternated a pong game and a tracking task (Fig. 2c). An experiment started with a **Track – Training** session that consisted of 30 trials (six for each target). The purpose of this session was to familiarize participants with the tracking task and to train them on the predictable figure-of-eight path. After training, participants were presented with a **Pong No Delay** session consisting of 10 trials. This was followed by a **Blind Track** session (**Post No Delay**) that consisted of 15 trials (three for each target). Next, participants experienced a **Pong Delay** session consisting of 30 trials. The time course of change in delay throughout the **Pong Delay** session in the Abrupt (N=10) and Gradual (N=10) groups was the same as in Experiment 2. The experiment ended with another **Blind Track** session (**Post Delay**) of 45 trials.

##### Experiment 4

In each experiment, sessions alternated a pong game and a tracking task (Fig. 2d). Each tracking session consisted of a single trial. An experiment started with a **Track – Training** session, followed by a **Blind Track – Training** session. The purpose of these sessions was to familiarize participants with the task. After training, participants were presented with a **Pong No Delay** session consisting of 10 trials. This was followed by a **Blind Track** session (**Post No Delay**). Next, participants experienced a **Pong Delay** session. The time course of change in delay throughout the **Pong Delay** session was the same as that of the Gradual groups in Experiments 2 and 3 for all participants (N=20). The experiment ended with another **Blind Track** session (**Post Delay**).

#### Simulations of the representation models

To control movements, it is commonly accepted that the brain performs state estimation of the body using sensory feedback (Wolpert et al., 1998; Shadmehr and Krakauer, 2008). Thus, to computationally formalize predictions of delay representation in the pong game, we assumed that the participants updated an estimate of the relationship between the hand location 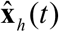 and the state of the visual feedback, the displayed paddle **x** _*p*_ (*t*). For the **Pong No Delay** session, we assumed that participants estimated the hand movement as being aligned with the movement of the paddle, and thus:

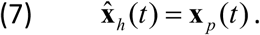

For the **Pong Delay** session, the hand moved according to the hand-paddle relationship that was predicted by each of the representation models. A *Time-based Representation* of the delay would lead to an estimate of hand location that explicitly included the actual time lag (*Τ*) between hand and paddle movements:

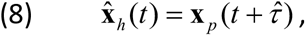

Where 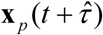 is the location of the paddle at estimated 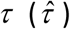 time ahead (Fig. 1c, left panel). A *State-based Representation* of the delay may follow one of three alternative models (Fig. 1c, right panel): participants may represent the current location of the hand according to the current location of the paddle spatially shifted by 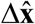 as a result of the delay (*Spatial Shift* – the paddle is constantly behind the hand):

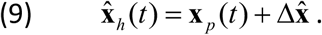

Alternatively, participants may attribute the distance between the hand and the delayed paddle to an altered proportional mapping 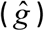 between the movement amplitudes of the hand and the paddle (*Gain* – the paddle moves in a smaller amplitude with respect to the amplitude of the hand):

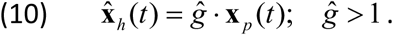

Another *State-based* alternative is to use a *Mechanical System* equivalent. A possible representation is that the paddle is a damped 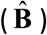 mass 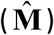 that is connected to the estimated representation of the hand position with a spring 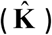

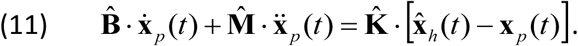

Such an approximation is based solely on the current state; i.e., the position, velocity and acceleration of the paddle. One possible choice of parameters in this representation can be calculated by considering a Taylor’s series approximation of the expression in Equation 8 around the position of the delayed paddle

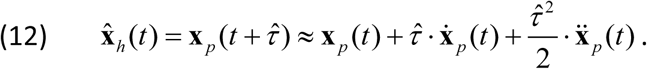

Rewriting Equation 11 as:

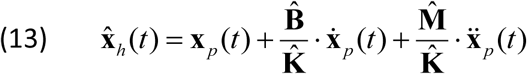

reveals that by choosing 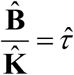 and 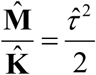 Equations 12 and 13 are equivalent.

We constructed predictions about the way each of the delay representation models affects the performance during the blind transfer tasks by simulating the predicted movements of each type of blind transfer task. Since during the transfer tasks participants were requested to imagine that there is a cursor, we assumed that they performed the task in the visual space, estimating the location of the imagined cursor 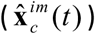 and attempting to place it in the target (which was either stationary during reaching or moving during tracking). Also, we assumed that the participants estimated 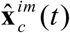 based on the position of the displayed paddle (X_p_(t)) during the former pong session such that 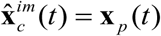. Thus, in our simulations, the hand moved according to the estimated relationship between the hand and the paddle. For the **Post No Delay** session, the simulated hand movement was based on a complete alignment between the hand and the paddle movements (Eq. 7). For the **Post Delay** session, the hand moved according to the hand-paddle relationship that was predicted by each of the representation models (Eqs. 8-10, 12).

##### Reaching

Reaching movements were simulated according to the minimum jerk trajectory (Flash and Hogan, 1985):

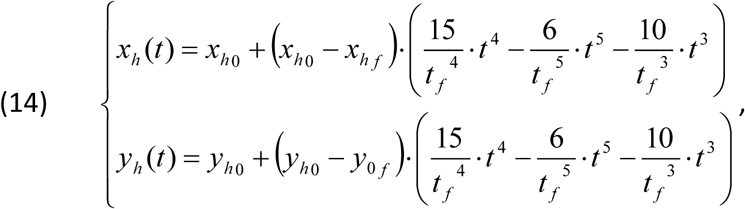

were *t _f_* = 0.3*s* was the movement duration; *x*_*h0*_ = *y*_*h0*_ = 0 *cm* and *xhf*, *yhf* were the initial and final hand positions coordinates of the simulated reaching movement, respectively. For the **Post No Delay** session, we set *x*_*hf*_ and *yhf* at the location of the targets in Experiment 1 such that the simulated movement amplitude was 10 cm. For the **Post Delay**, we simulated the predicted hand trajectories according to each of the representation models such that the imagined cursor / paddle would reach the target (Eqs. 8-10, 12). We chose the parameters that, when possible, produced the effects that were similar in magnitude to the effects that were observed in the experiment. For *Time Representation* of the delay, we presented the simulation results for 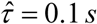 (Eq. 8). For the *State Representation* models, the reaching movements for the *Spatial Shift* model were generated with the free parameter 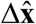 (Eq. 9) equals to 1.5*cm*; the results for the *Gain* model were generated with the free parameter 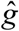 (Eq. 10) equals to 1.2; and the results for the *Mechanical System* model were generated with a free parameter of the Taylor’s series approximation 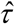 (Eq. 12) equals to 0.1 s.

To present the simulation results (Fig. 4c) in a consistent manner with the presentation of the experimental results, we added noise to the endpoint of each simulated movement. The noise was drawn from a normal distribution with zero mean and 1 cm standard deviation. This noise is thought to correspond to the noise present in various stages of sensorimotor control (Franklin and Wolpert, 2011).

**Fig 4.**
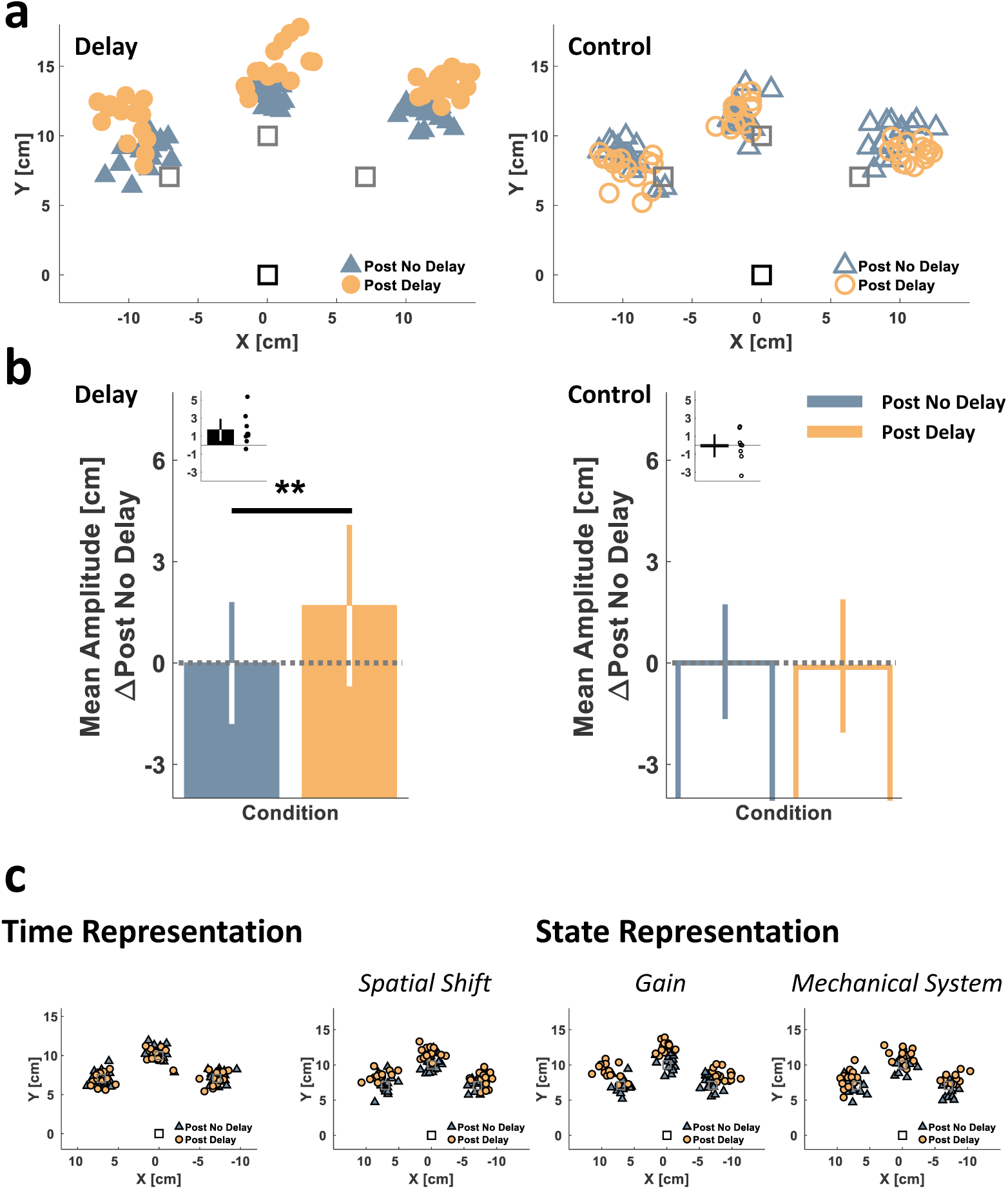
Experiment 1: reaching experimental results and representation model simulation results suggest a *State-based* rather than a *Time-based Representation* of delay. (a) Single participant’s experimental results from each of the Delay (left, filled markers) and Control (right, hollow markers) groups. Movements start location is indicated by the black square and target locations are marked by the gray squares. Markers represent the end point locations of the hand at movement terminations during the **Post No Delay** (blue triangles) and **Post Delay** (orange circles) **Blind Reach** sessions. (b) Experimental results group analysis. Colored bars represent the mean reaching movement amplitudes towards all targets of each participant, and for each of the **Blind Reach** sessions, averaged over all the participants in each group (Delay: left, N=9, Control: right, N=8) and following subtraction of each group’s average baseline amplitude (during the **Blind Reach – Post No Delay** session). Black bars (insets) represent the difference in mean amplitude between the **Post Delay** and the **Post No Delay** blind reaching sessions for each participant, averaged over all targets and over all the participants in each group. Error bars represent the 95% confidence interval. (c) Simulation results of reaching end points in the Delay group (**Post No Delay** – black outlined blue triangles, **Post Delay** – black outlined orange circles) for *Time Representation* (left) and *State Representation* (right) of the delay. **p<0.01.

##### Tracking – figure-of-eight

Tracking movements were simulated for a complete single cycle of the target movement along the sagittal dimension of the figure-of-eight path (Fig. 7). Thus, each hand trajectory was simulated as a single sine cycle. Since accurate performance during such a task is very rare, participants may exhibit various tracking errors even during baseline. However, the predicted relative effects of the delay are valid regardless of baseline accuracy. Thus, for illustration purposes, we assume that during the **Post No Delay** session, the hand lagged behind the movement of the target (Rohde et al., 2014) by 0.2 s. For the **Post Delay** session, we simulated the effect of each delay representation model on the resulting hand movement. For *Time Representation*, we presented the simulation results for 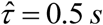 (Eq. 8) (0.7 s relative to baseline). For the *State Representation* models, the tracking movements for the *Spatial Shift* model were generated with 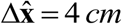 (Eq. 9); those for the *Gain* model were generated with 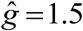 (Eq. 10); and the results for the *Mechanical System* model were generated with the values of 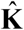, 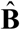 and 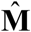 that fulfill 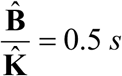 and 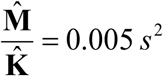 (Eq. 13). Note that we did not draw any conclusions from the magnitudes of the parameters’ in the representations; we chose parameters that resulted in an observable change in the hand trajectory due to the delay and that could illustrate the effects qualitatively.

**Fig 7.**
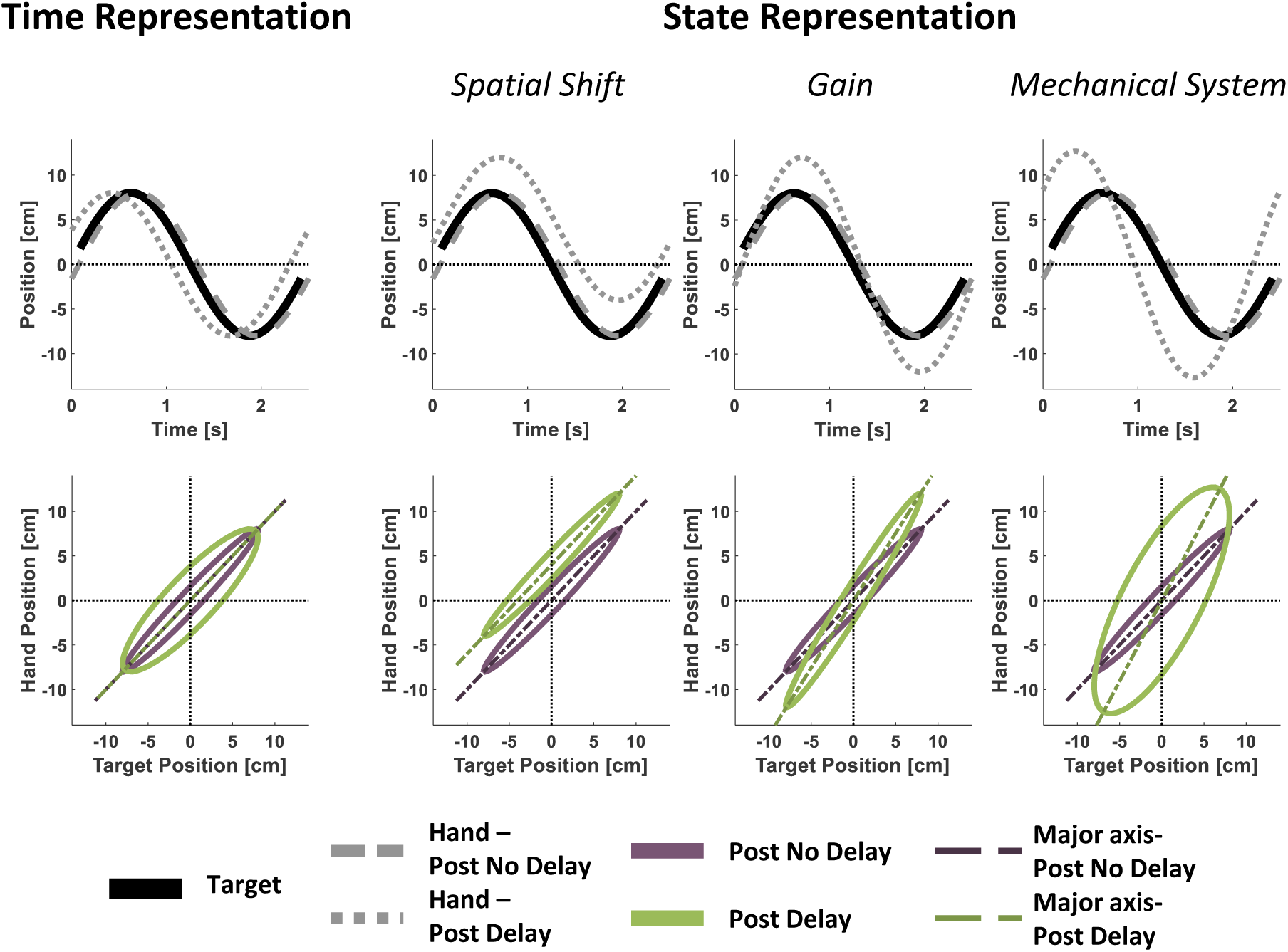
Experiment 3: blind tracking predictions.

##### Tracking – mixture of sinusoids

We simulated frequency responses for tracking movements in different frequencies to illustrate the predicted effect of delay representation as *Gain* and *Mechanical System* on frequency-dependent increase in movement amplitude (Fig. 9). The simulation was conducted for a target movement that had an amplitude of *A*_*t*_ = 2 *cm* for all movement frequencies (*fr*) within the range of [0 1]. For an accurate baseline (**Post No Delay**) tracking performance, we simulated hand amplitude (*A*_*h*_) that was equivalent to the target amplitude at all movement frequencies: *A*_*h*_ = *A*_*t*_. For the **Post Delay** session, we calculated for the *Gain* model the predicted hand amplitude in all frequencies with 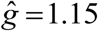 For the *Mechanical System* model, we used the Fourier transform of Equation 12 (for the sagittal plane, which was the only dimension in which the target was moving), and calculated the transfer function of the hand-paddle relationship:

**Fig 9.**
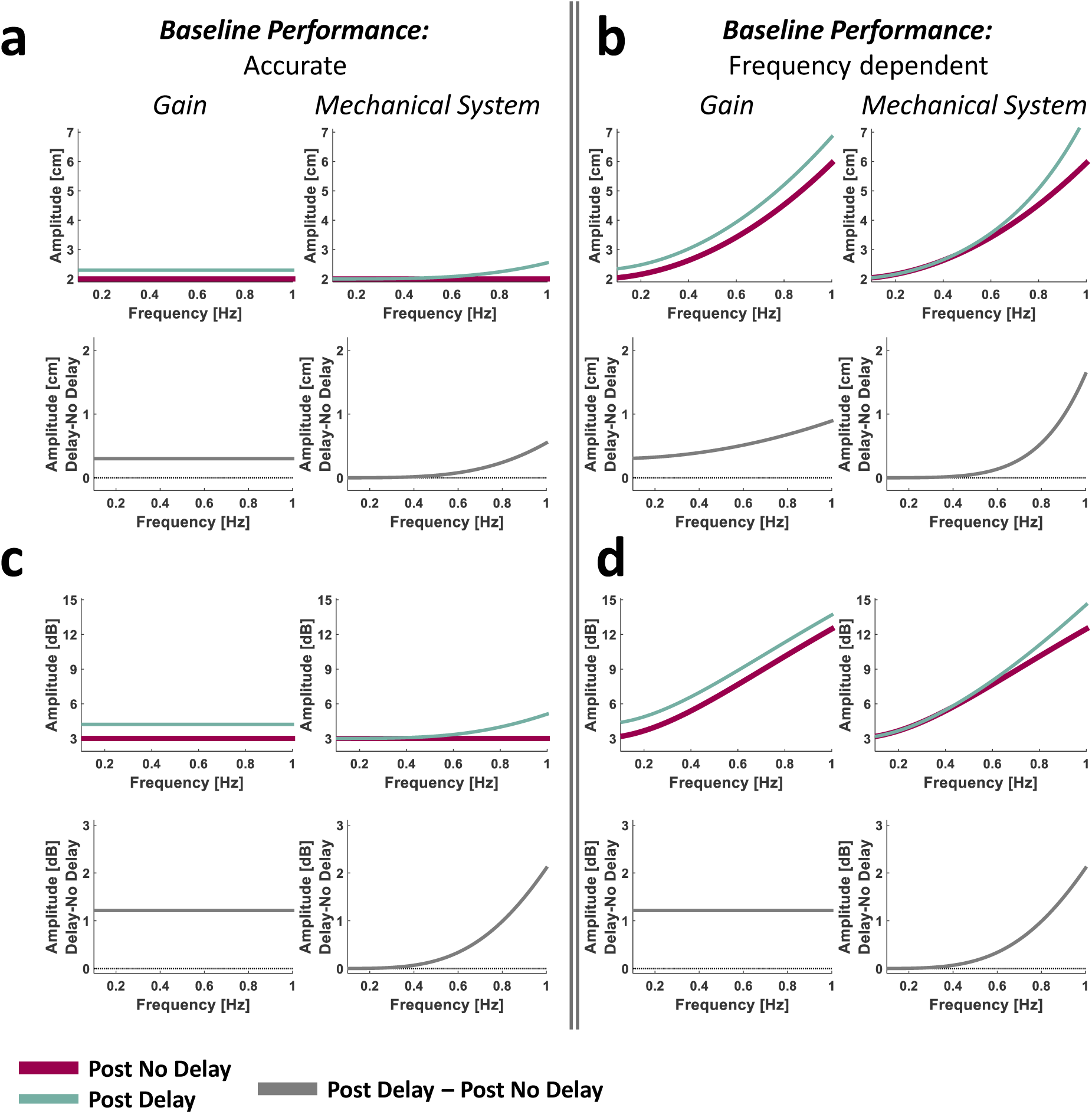
Experiment 4: predicted frequency effects on delay-induced hypermetria.

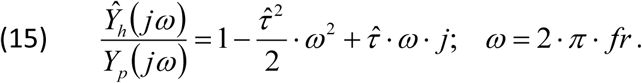

Thus, the predicted hand amplitude for a target moving at an amplitude of *A*_*t*_ with a *Mechanical System* representation is:

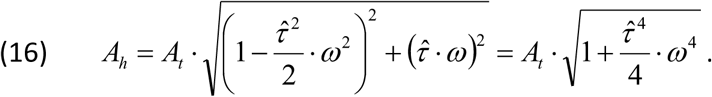

The simulation results for this model were generated with 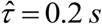 frequency responses for both the amplitude in metric scale (*A*_*h*_, in cm) and the decibel amplitude (*DA*_*h*_, in dB). We calculated the latter as *DA*_*h*_ = 10 ⋅ log _10_ (*pow*), where 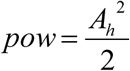 is the power associated with each movement frequency.

To illustrate the effect of baseline accuracy on the predicted amplitude for each model, we also simulated the frequency responses for the case of an increase in the baseline movement amplitude with an increase in the movement frequency (Foulkes and Miall, 2000). We presented the simulation for the function *A*_*h*_ = *A*_*t*_ + 0.1⋅ *ω* 2, which exhibits an increase in the examined range of *ω*.

##### Pong

We simulated frequency responses for the movements during the pong game to illustrate the predicted effect of all the representation models on changes in movement amplitudes due to the delay (Fig. 11c). To conduct the simulation, we averaged the frequency response profiles of the last four trials of the **Pong No Delay** session from a representative participant (Fig. 11b, black) (see Data analysis – Metrics); we used this mean profile as an example for a baseline frequency response in the pong game. For the **Delay** session, we used the same frequency response mean profile for both the *Time* and *Spatial Shift* models as they are not associated with any change in the movement amplitude. For the *Gain* model, we calculated the predicted hand amplitude in all frequencies with 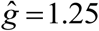 For the *Mechanical System* model, we used the same transfer function of the hand-paddle relationship as with the tracking – mixture of sinusoid simulation (Eq. 15) and calculated the predicted hand amplitude using Equation 16 (this time, in Eq. 16 is the baseline frequency response profile). The simulation results for this model were generated with 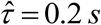. We presented the predicted frequency responses for the amplitude in metric scale and the frequency response difference profile between the **Delay** and **No Delay** sessions in decibel units.

**Fig 11.**
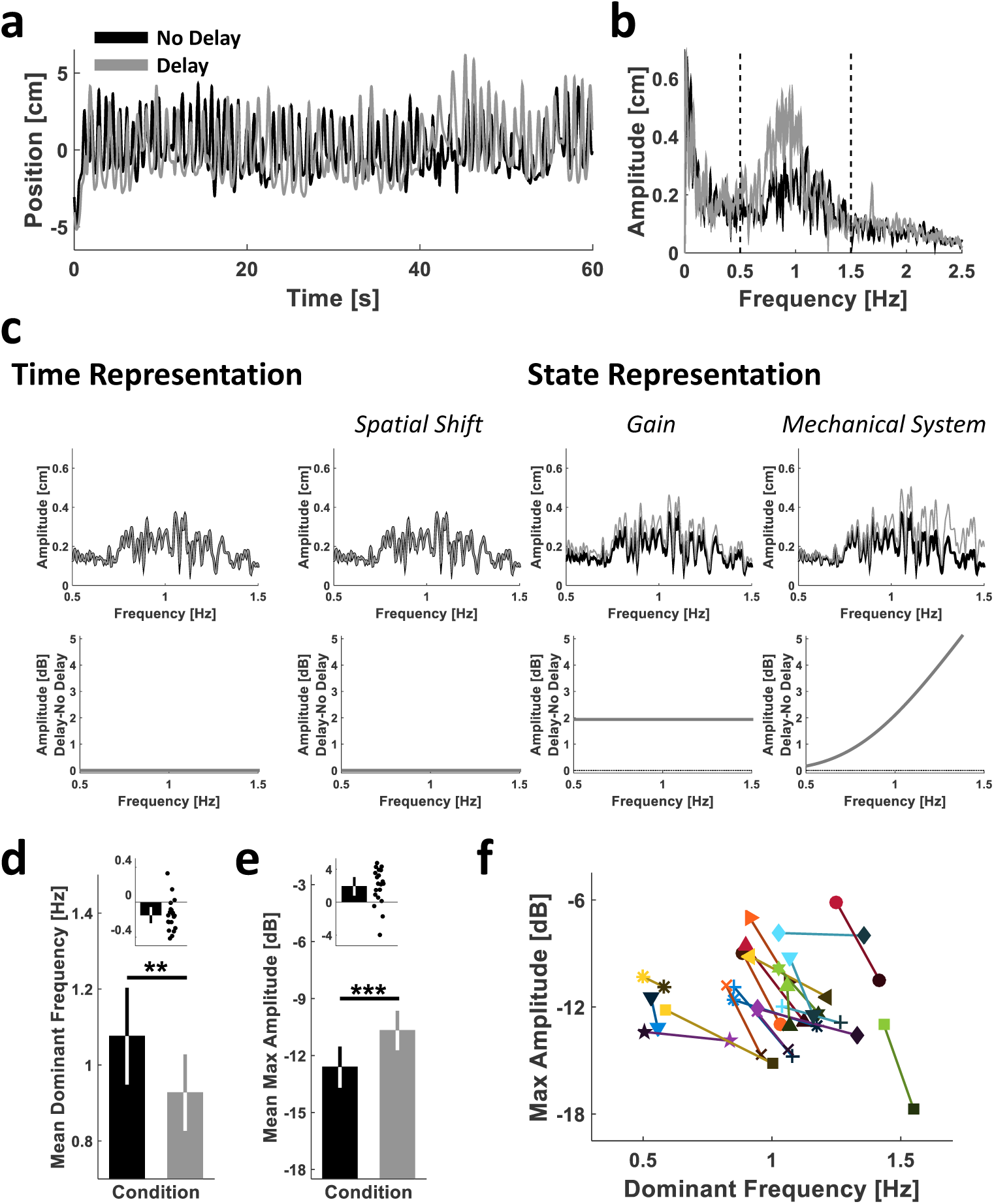
Experiment 4: frequency response analysis of pong movements and representation model simulation results are most consistent with the *Gain* representation model. (a, b) Single participant’s results. Sagittal hand trajectories of a representative participant during the last pong trial of each of the **No Delay** (black) and **Delay** (gray) sessions (a), and the mean frequency responses of the sagittal hand trajectories from the last four trials of each session (b). The vertical dashed lines define the frequency range of interest within which the participants were mainly moving ([0.5 1.5] Hz). (c) Simulation results of the predicted effect of delay according to each of the representation models, illustrated using the baseline (no delay) frequency response of the participant in (b). Upper panels display the **No Delay** (black) and the **Delay** (gray) amplitudes in cm, and lower panels present the difference between the amplitudes in dB. (d-e) Group analysis. Mean dominant frequency (d) and mean maximum amplitude (e) of all participants (N=20). The black bars (inset) represent the mean difference in each measure between the **Delay** and the **No Delay** pong sessions. Error bars represent the 95% confidence interval. Dots represent differences of individual participants. (f) The maximum amplitude and its respective frequency (dominant frequency) for each participant is presented in a frequency-amplitude space to illustrate the overall changes dynamic of both measures from the **No Delay** (dark markers) to the **Delay** (light markers) pong session. **p<0.01; ***p=0.001.

#### Data analysis

### Metrics

Device position, velocity, and the forces applied were recorded throughout the experiments at 200 Hz. They were analyzed off-line using custom-written MATLAB^®^ code (The MathWorks^®^, Inc., Natick, MA, USA, RRID: SCR_001622).

#### Pong: Hit rate

To examine performance in the pong game, we analyzed the change in the paddle-ball hit rate throughout the experiment. As mentioned above, the ball changed its movement direction from down to up when it either hit the bottom wall of the arena or during a hit. Thus, we identified the number of hits off-line by extracting the number of times the ball movement direction changed upward and its sagittal location was not at the bottom wall at the time of the change. Since the duration of each of the Pong sessions in Experiment 1 varied across participants, to analyze the changes in the average hit rate of all participants in each group, for each participant, we pooled the data of a session and divided it into bins of equal duration. The **Pong No Delay** session was divided into five bins, and the **Pong Delay** session was divided into 20 bins. Hit rate was calculated as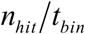, where *nhit* was the number of hits in a bin. In Experiment 2, the duration of the Pong sessions was equal between participants, and consisted of the same number of trials, each with a duration of *t*_*Trial*_ = 60 *s*. Thus, in these experiments, hit rate was calculated as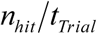, where *n*_*hit*_ was the number of hits in a trial.

#### Reaching: Amplitude

For the purpose of data analysis, we defined movement onset at the first time the velocity exceeded two percent of its maximum value. Movement end time was set at 0.1 s after the velocity dropped below five percent of its maximum value; the reaching end-point was thus defined as the hand location (**x**_*h*_) at that time point. Reaching amplitude was calculated as the Euclidean distance between **x**_*h*_ at movement onset and movement end-point.

#### Tracking – figure-of-eight: Target-Hand Delay, Slope, and Intercept (Experiment 3)

As mentioned above, during each figure-of-eight tracking trial, the tracking task began immediately after the participant reached towards a target within the figure-of-eight path and stopped. Thus, we segregated the tracking movement from the reaching movement by defining tracking onset as the first sampled time point in which the target started moving.

To evaluate tracking accuracy, we calculated an *R*^2^ value for each tracking trial according to (Nagengast et al., 2009):

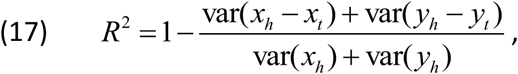

where var was the variance of the expression in parentheses. In 12% of the individual **Blind Track** trials, the *R*^2^ was less than 0.6, and were omitted from further analyses.

Since the pong game was two dimensional, we analyzed the effect of the game on both the *x*_*h*_ (*t*) and *y*_*h*_ (*t*) components of the hand movement that tracked the two-dimensional target path (Eq. 5). To measure Target-Hand Delay, for each dimension, we calculated the cross correlation between target and hand positions (*x*_*t*_ (*t*) and *x*_*h*_ (*t*) for the frontal dimension, and *y*_*t*_ (*t*) and *y*_*h*_ (*t*) for the sagittal dimension) on each trial, and found the lag for which the cross correlation was maximal. Positive values of Target-Hand Delay indicate that the hand movement preceded the movement of the target. The purpose of this measure was to examine whether participants used a *Time-based Representation* to cope with the delay. If they did, the predicted effect would be an increase in the Target-Hand Delay from the **Post No Delay** to the **Post Delay** tracking session.

As illustrated in Figures 7 and 8, we examined the relationship between the target and the hand during tracking by projecting the sampled position of each in a target-hand position space. Then, we fit an ellipse to the data points with the following form (Fitzgibbon et al., 1999; Chernov, 2009):

**Fig 8.**
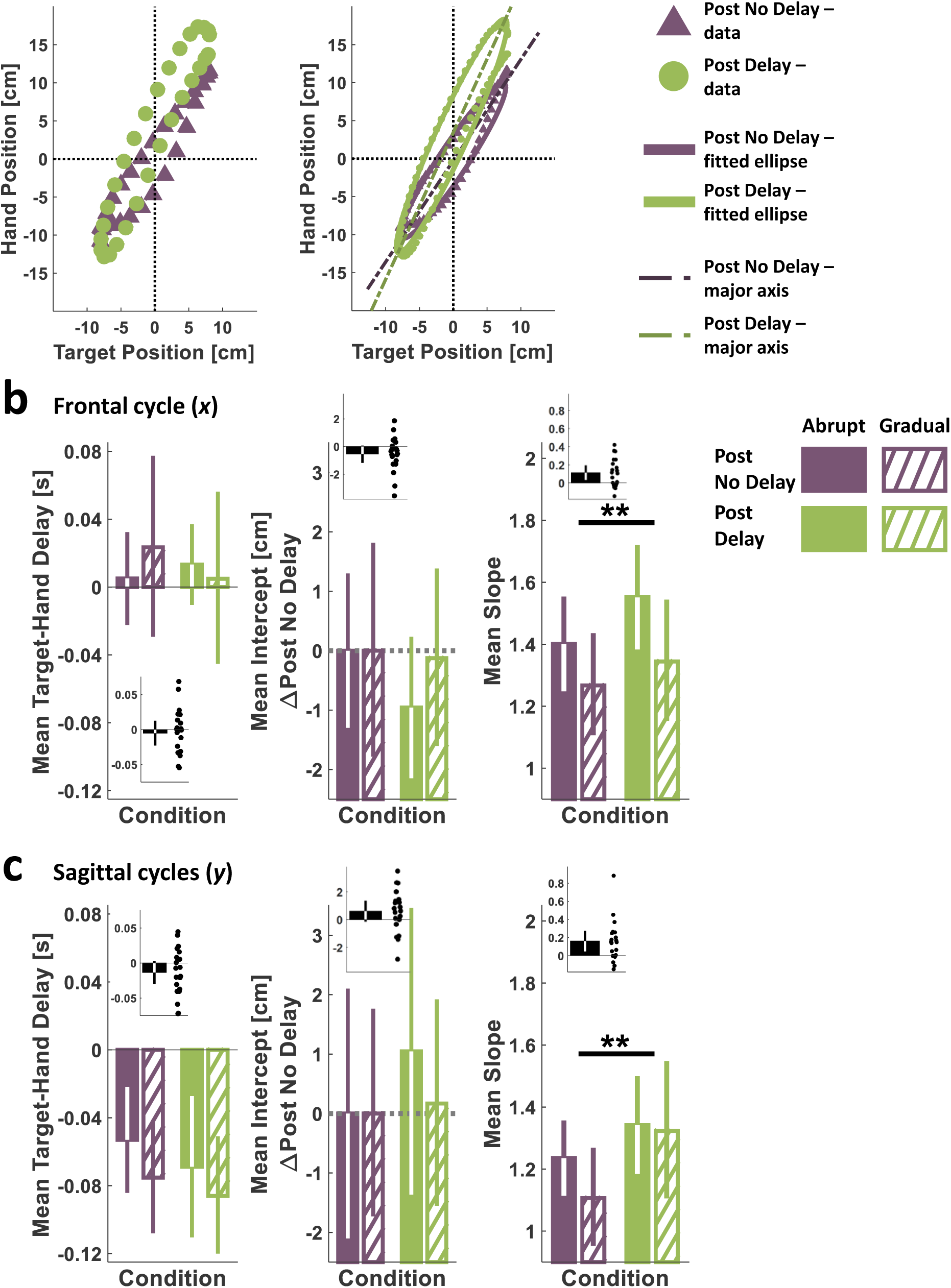
Experiment 3: tracking experimental results suggest a *State Representation* of delay as either a *Gain* or a *Mechanical System* equivalent rather than a *Spatial Shift*. (a) Single participant’s results. Target-hand position space of a single sagittal cycle from each of the **Post No Delay** (purple triangle) and **Post Delay** (green circles) **Blind Track** sessions. The left panel presents data points sampled at 11.8 Hz. The right panel presents data points sampled at 28.6 Hz and the fitted ellipses for entire data distribution (sampled at 200 Hz) from each of the **Post No Delay** (purple) and **Post Delay** (green) tracking sessions, together with the corresponding major axis lines (dashed-dotted dark purple and dashed-dotted dark green, respectively). (b,c) Group analyses for the frontal cycle (b) and for the sagittal cycles (c) of the delay between the hand and the target (left), and the major axis intercepts (after subtraction of each group’s average **Post No Delay** intercept, middle) and slopes (right), extracted from participants’ tracking performances. Colored bars represent each participant’s mean, from each of the **Post No Delay** (purple) and **Post Delay** (green) tracking sessions, averaged over all the participants in each group (Abrupt: filled, N=10, Gradual: diagonal lines, N=10). The black bars (insets) represent the mean difference for each measure between the **Post Delay** and the **Post No Delay** blind tracking sessions. **p<0.01.

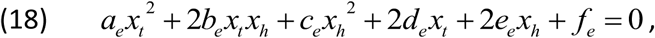

where *x*_*t*_ and *x*_*h*_ are the Euclidean space coordinates of the target and hand frontal movement direction in a single trial, respectively. The same was done also for *y*_*t*_ and *y*_*h*_ – the Euclidean space coordinates of the target and hand sagittal movement direction. Note that the figure-of-eight is constructed from a single frontal sine cycle and two sagittal cycles, but for each dimension we fitted a single ellipse for all the data points. Then, we extracted the Slope and the Intercept of the ellipse’s major line. To do this, we derived the coordinates of the center of the ellipse (*o*_*t*_, *o*_*h*_) according to:

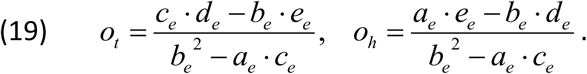

The counterclockwise angle of rotation (*θ*) between the *x*_*t*_ or the *y*_*t*_ axis and the ellipse’s major line is:

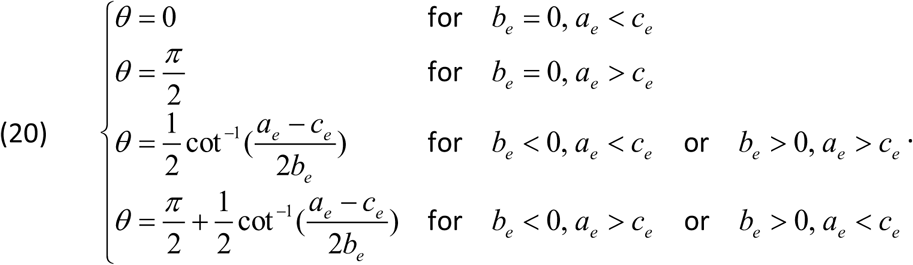

The ellipse’s major line Slope (*s*_*maj*_) and Intercept (*i*_*maj*_) were calculated according to:

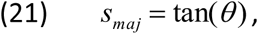

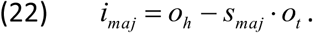

The Slope and Intercept measures were used to assess how the *State Representation* of the delay takes place; an increase in the Intercept suggests a representation of delay in the form of a *Spatial Shift*, whereas an increase in the Slope is consistent with a delay representation as a *Gain* or a *Mechanical System*.

#### Tracking – mixture of sinusoids: Frequency response (Experiment 4)

To measure the hand amplitude for each of the main frequencies in the tracking movement, we calculated the periodogram power estimate for each hand trajectory using the Matlab function *periodogram()* and with a Hanning window (Matlab’s *hann()* function). To obtain accurate estimates of the amplitudes in the sharp peaks of the discrete Fourier transform in our experiment, each hand trajectory vector (∼24000 samples length) was padded with zeros to a vector length of 600,000 samples. Then, we extracted the five peak power estimates associated with each of the five frequencies in the target trajectory. The amplitude (*A*_*h*_, in cm, Fig. 10b) was calculated from the power (*pow*) as 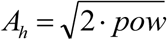 To examine the effect of delay, we calculated the decibel amplitude (*DA*_*h*_, in dB units, Fig. 9c) from the power as *DA*_*h*_ = 10 ⋅ log _10_ (*pow*). Finally, we calculated the difference in DA between the **Post Delay** and the **Post No Delay** sessions (Fig. 10d).

**Fig 10.**
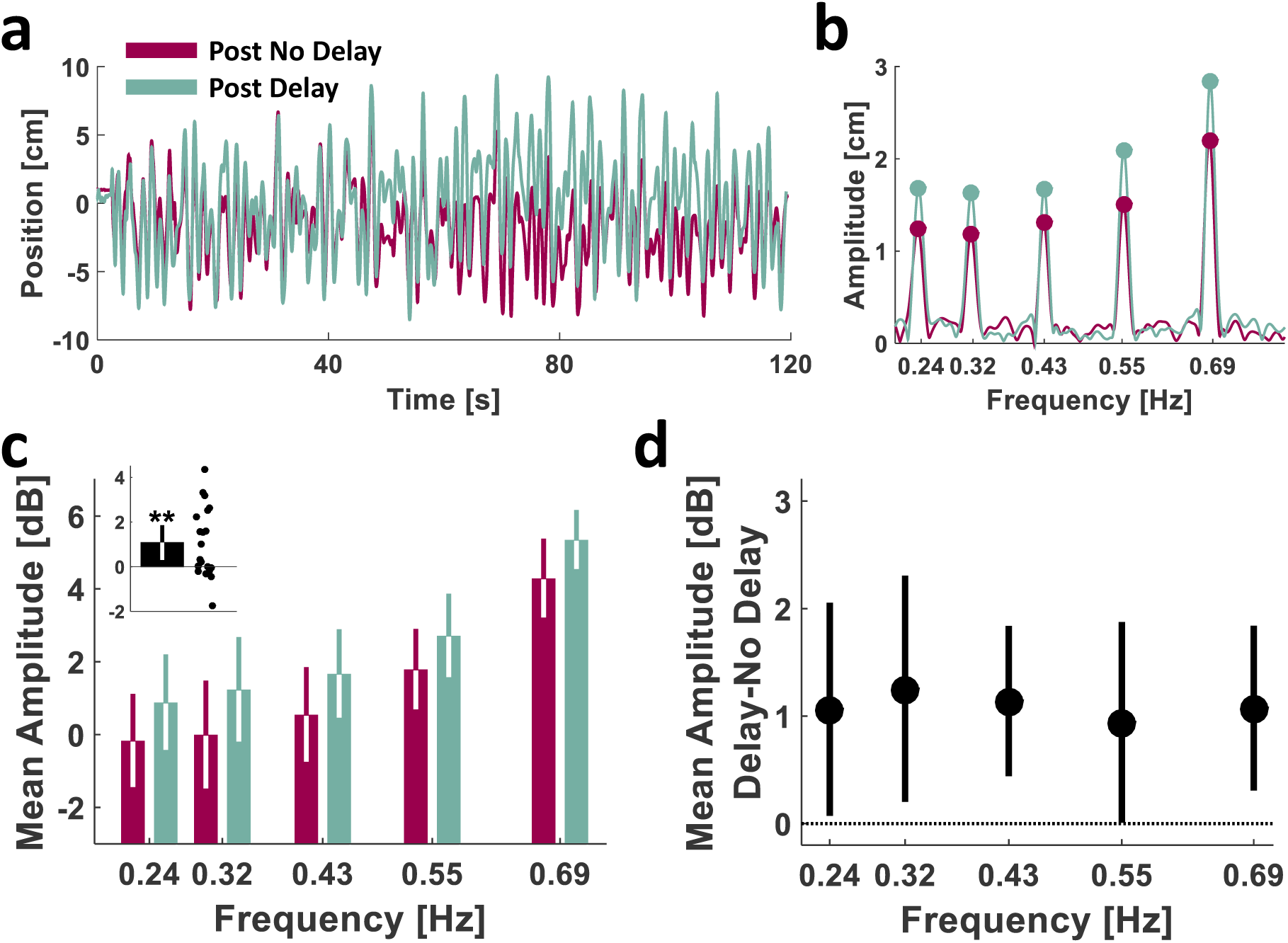
Experiment 4: experimental results for tracking with different frequencies suggest a.

#### Pong: Frequency response (Experiment 4)

To measure the change in hand amplitude due to the delay during the pong game, we calculated the Fast Fourier Transform (*FFT*) for each hand trajectory from the last four trials of each of the **Pong No Delay** and the **Pong Delay** sessions using the Matlab function *fft()*. Since the design of our pong game encouraged participants to repetitively hit the ball towards the upper wall of the arena, we focused our analysis on the sagittal component of the hand movement. Prior to the FFT calculation, each hand trajectory vector (∼12000 samples length) was padded with zeros to a vector length of 300,000 samples. For each trajectory, we calculated the amplitude as 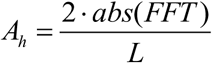, where *L* is the length of the original hand trajectory vector (prior to the zero padding), and the decibel amplitude as 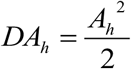 for all movement frequencies. Then, for each participant, we averaged the frequency responses of the four trials in each stage. Visual examination of the responses revealed that participants were mainly moving within the [0.5 1.5] Hz frequency range, and therefore, we focused on the responses within this range (we also observed a low frequency (<0.2 Hz) peak that is due to pauses during the game, and is less interesting in terms of dynamic delay perturbation). For each of the mean *DA*_*h*_ frequency responses, we filtered the mean responses by calculating the centered moving average with a window size of 101 samples and found the maximum decibel amplitude and its corresponding frequency.

### Statistical analysis

Statistical analyses were performed using custom written Matlab functions, Matlab Statistics Toolbox, and IBM^®^ SPSS (RRID: SCR_002865). The raw data and custom software will be made available upon publication.

We used the Lilliefors test to determine whether our measurements were normally distributed (Lilliefors, 1967). For ANOVA models that included a within-participants independent factor with more than two levels, we used Mauchly’s test to examine whether the assumption of sphericity was met. When it was not, F-test degrees of freedom were corrected using the Greenhouse-Geisser adjustment for violation of sphericity. We denote the p values that were calculated using these adjusted degrees of freedom as *p*_*ε*_. For the factors that were statistically significant, we performed planned comparisons, and corrected for family-wise error using a Bonferroni correction. We denote the Bonferroni-corrected p values as *p*_*B*_.

In Experiment 1, to analyze the change in hit rate throughout the experiment for each of the Delay and Control groups, for each participant we calculated the mean hit rate of the last four bins in the **Pong No Delay** session (Late No Delay), and the first (Early Delay) and last (Late Delay) four bins in the **Pong Delay** session. Then, we fit a two-way mixed effect ANOVA model, with the mean hit rate as the dependent variable, one between-participants independent factor (Group: two levels, Delay and Control), and one within-participants independent factor (Stage: three levels, Late No Delay, Early Delay and Late Delay).

In Experiment 2, to analyze the change in hit rate throughout the **Pong Delay** session and to compare the Abrupt and Gradual groups, for each participant we calculated the mean hit rate of the last five trials in the **Pong No Delay** session (Late No Delay), and the first (Early Delay) and last (Late Delay) five trials in the **Pong Delay** session. Then, we fit a two-way mixed effect ANOVA model, with the mean hit rate as the dependent variable, one between-participants independent factor (Group: two levels, Abrupt and Gradual), and one within-participants independent factor (Stage: three levels, Late No Delay, Early Delay and Late Delay).

To analyze the effect of the delayed pong on reaching amplitude, for each participant we evaluated the mean reaching amplitude during the **Post No Delay** and **Post Delay** sessions. We fit a three-way mixed effects ANOVA model, with the mean reaching amplitude as the dependent variable, one between-participants independent factor (Group: two levels, Experiment 1: Delay and Control, Experiment 2: Abrupt and Gradual), and two within-participants independent factor (Session: two levels, Post No Delay and Post Delay. Target: three levels, Right, Middle and Left). Mauchly’s test indicated a violation of the assumption of sphericity for the main effect of Target on the reaching amplitude in Experiment 2 (*χ*^2^ (2) = 6.507, *p* = 0.039), and for the Session and Target interaction effect (*χ*^2^ (2) = 12.028, *p* = 0.002). Thus, we applied the Greenhouse-Geisser correction factor to the Target factor’s degrees of freedom in the former 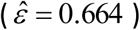, and to the Session-Target and the Session-Target-Group interactions’ degrees of freedom in the latter 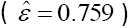.

To analyze the effect of the delayed Pong on the figure-of-eight tracking performance in Experiment 3, for each participant we evaluated the mean Target-Hand Delay, Slope and Intercept measures for each movement dimension during the **Post No Delay** and **Post Delay** sessions. For each measure, we fit a two-way mixed effect ANOVA model, with the measure as the dependent variable, one between-participants independent factor (Group: two levels, Abrupt and Gradual), and one within-participants independent factor (Session: two levels, Post No Delay and Post Delay).

To analyze the effect of the delayed Pong on the mixture of sinusoids tracking performance in Experiment 4, for each participant we evaluated the decibel amplitude of the main five movement frequencies during the **Post No Delay** and **Post Delay** sessions. We fit a two-way repeated measures ANOVA model, with the decibel amplitude as the dependent variable, and two within-participants independent factors (Session: two levels, Post No Delay and Post Delay. Frequency: five levels). Mauchly’s test indicated a violation of the assumption of sphericity for the main effect of Frequency on the tracking amplitude (*χ*^2^ (9) = 30.383, *p* < 0.001), and for the Session-Frequency interaction effect (*χ*^2^ (9) = 25.066, *p* = 0.003).

We used a two-tailed *paired-sample t*-test to examine the effect of the delayed pong on the Target-Hand delay in the tracking task of Experiment 4, and on the maximum decibel movement amplitude and its corresponding frequency (dominant frequency) in the pong game.

Throughout this paper, statistical significance was set at the *p* < 0.05 threshold.

## Results

### Experiments 1 & 2

#### Transfer of hypermetria following a delayed pong game to a blind reaching task suggests State rather than Time Representation of the delay

In Experiment 1, the Delay (N=9) and the Control (N=8) groups played two **Pong** sessions (Fig. 2a). To evaluate performance in the pong game, we calculated the paddle-ball hit rate and analyzed its change throughout the experiment in both the Delay and Control groups (Fig. 3). The change in hit rate throughout the stages of the experiment was different between the groups (Stage-Group interaction effect: *F*_(2,30)_ = 20.512, *p* < 0.001). The hit rate of the Control group, who did not experience a delay in both **Pong** sessions, remained the same throughout the experiment (Late No Delay – Early Delay: *p*_*B*_ = 0.982; Late No Delay – Late Delay: *p*_*B*_ =1.000; Early Delay – Late Delay: *p*_*B*_ = 0.438). However, as a result of the sudden presentation of the delay, the hit rate of the Delay group decreased drastically (*p*_*B*_ < 0.001), and then increased with continued exposure to delay (*p*_*B*_ = 0.023). Yet, they did not reach the same hit rate as during Late No Delay (*p*_*B*_ < 0.001) or the Control group at the corresponding Late Delay stage (*p*_*B*_ = 0.004). Thus, participants from the Delay group were able to improve their performance during exposure to the delay, but this improvement was mild, suggesting a difficulty in adapting to the perturbation.

**Fig 3.**
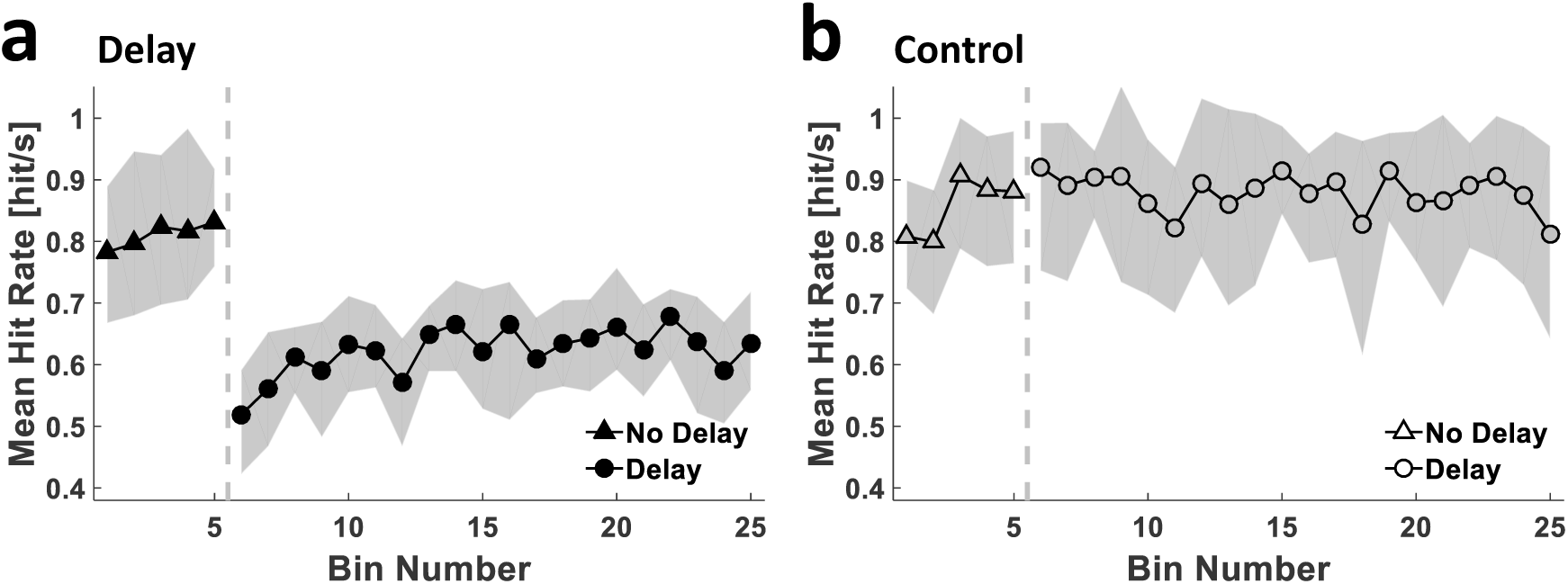
Experiment 1: paddle-ball hit rate in the presence of delayed and non-delayed feedback.

Time courses of the mean hit rate of all participants in each of the Delay (a, filled markers, N=9) and Control (b, hollow markers, N=8) groups. The grey dashed vertical line separates the **Pong No Delay** (triangles) and the **Pong Delay** (circles) sessions. Shading represents the 95% confidence interval.

Both the Delay and the Control groups performed sessions of a blind reaching task after the two **Pong** sessions (Fig. 2a, blue and orange frames). This enabled us to capture the representation of hand-paddle dynamics following exposure to either the non-delayed or the delayed pong game where participants had to rely solely on a feedforward mechanism and proprioceptive feedback. Analysis of participants’ performance in the blind reaching task revealed that participants from the Delay group, but not the Control group, made longer (hypermetric) reaching movements after the delayed pong game. Figure 4a presents the reaching endpoints – the locations of movement terminations – during the **Post No Delay** and **Post Delay** blind reaching sessions from a representative participant in each group. Whereas for the participant in the Delay group, **Post Delay** movement endpoints reached farther from the start location than the **Post No Delay** movements’ endpoints (Fig. 4a, left), for the participant in the Control group, the blind reaching movements from the **Post No Delay** and **Post Delay** sessions ended at around the same location (Fig. 4a, right).

We extracted the reaching amplitude from all movements in each session (Fig.4b). Playing pong in the presence of a delay affected reaching amplitudes (Session-Group interaction effect: *F*_(1,15)_ = 4.717, *p* = 0.046). For participants in the Delay group, the reaching amplitude significantly increased from the **Post No Delay** to the **Post Delay** session (Post Delay – Post No Delay: [*mean difference, 95% CI*], 1.697 *cm*, [0.470 2.925], *p*_*B*_ = 0.010) (Fig 4b, left). A similar increase was not found in the Control group (− 0.126 *cm*, [−1.427 1.176], *p*_*B*_ = 0.840) (Fig.4b, right). Overall, these statistical analyses suggest that the specific experience with the delayed pong caused the participants to perform larger blind reaching movements.

Analysis showed that participants made larger movements towards the right target than they did towards the other targets (main effect of Target: *F*_(2,30)_ = 59.581, *p* < 0.001). For both Delay and Control groups, the reaching amplitudes to the right target were larger than to the left (*p*_*B*_ < 0.001) and to the middle (*p*_*B*_ < 0.001) targets. In addition, for the right target alone (Target-Session interaction effect: *F*_(2,30)_ = 10.175, *p* < 0.001), there was a statistically significant increase in movement amplitude between the **Post No Delay** and the **Post Delay** blind reaching sessions (*p*_*B*_ = 0.007). No such differences were found for the left (*p*_*B*_ = 0.808) and for the middle targets (*p*_*B*_ = 0.167). Importantly, these differences in reaching amplitudes between the targets did not stem from the applied delay (Group-Target-Session interaction effect: *F*_(2,30)_ = 0.299, *p* = 0.744). Thus, we reasoned that they stemmed from biomechanical differences in reaching towards different directions (Mussa-Ivaldi et al., 1985; Carey et al., 1996), from the difficulty of reaching to visual targets without visual feedback of the hand, and potentially, insufficient training on this task. Therefore, in Experiment 2, we added an additional session at the beginning of the experiment to train the participants on the blind reaching task.

To understand which of the representation models depicted in Figure 1c best accounted for the observed results, we simulated reaching movements towards targets for the **Post No Delay** and **Post Delay** conditions of the Delay group based on four models: *Time Representation, State Representation – Spatial Shift, State Representation – Gain* and *State Representation – Mechanical System*.

The simulation results are presented in Figure 4c. The **Post No Delay** endpoints were closely distributed around the target locations. For *Time Representation* of the delay, in which an estimate of the actual time delay was available (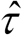 in Eq. 8), the **Post Delay** endpoints were also distributed around the target locations, and were not influenced by the value chosen for the estimated delay parameter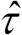. Hence, there was no parameter value in the *Time Representation* model yielding simulation results that were consistent with the reaching overshoot observed in the experimental results. In contrast, for all the *State Representation* models (the *Spatial Shift*, the *Gain* and the *Mechanical System*), we identified parameter values which resulted in simulated **Post Delay** overshoots similar to the experimental observations. Thus, *State Representation* and not *Time Representation* appeared to be able to account for the increase in movement amplitude following the delayed pong task.

#### Hypermetria is comparable in the abrupt and gradual conditions

The group that experienced the delay in Experiment 1 which then exhibited hypermetric movements during transfer to a blind reaching task was presented with an abrupt delay perturbation. Since adaptation through gradually increasing perturbations was shown to enhance transfer (Kluzik et al., 2008; Torres-Oviedo and Bastian, 2012), we hypothesized that presenting participants with a gradually increasing delay during the **Pong Delay** session would result in an increase in the reaching movement amplitude during the blind transfer task compared to the abrupt case. To test this hypothesis, we ran a second experiment (Experiment 2) in which we compared between a gradual (Gradual, N=10) and abrupt (Abrupt, N=10) presentation of the delay.

The analysis of the paddle ball hit rate (Fig. 5) revealed that the change in the hit rate throughout the delayed pong session differed between groups (Group-Stage interaction effect: *F*_(2,36)_ = 18.546, *p* < 0.001). Participants in the Abrupt group improved their performance in the presence of the delay (*p*_*B*_ = 0.006). In contrast, since the Gradual group did not experience an abrupt change in the delay, the mean hit rate of these participants was higher than that of the Abrupt group at the beginning of the **Pong Delay** session (*p*_*B*_ < 0.001). As the delay increased, there was a decrease in their performance (*p*_*B*_ = 0.001). Altogether, while these results suggest that the Abrupt group adapted to the delay, due to the increase in the delay in the Gradual protocol, which may conceal a possible tendency towards improvement, we cannot claim the same for the participants in the Gradual group.

**Fig 5.**
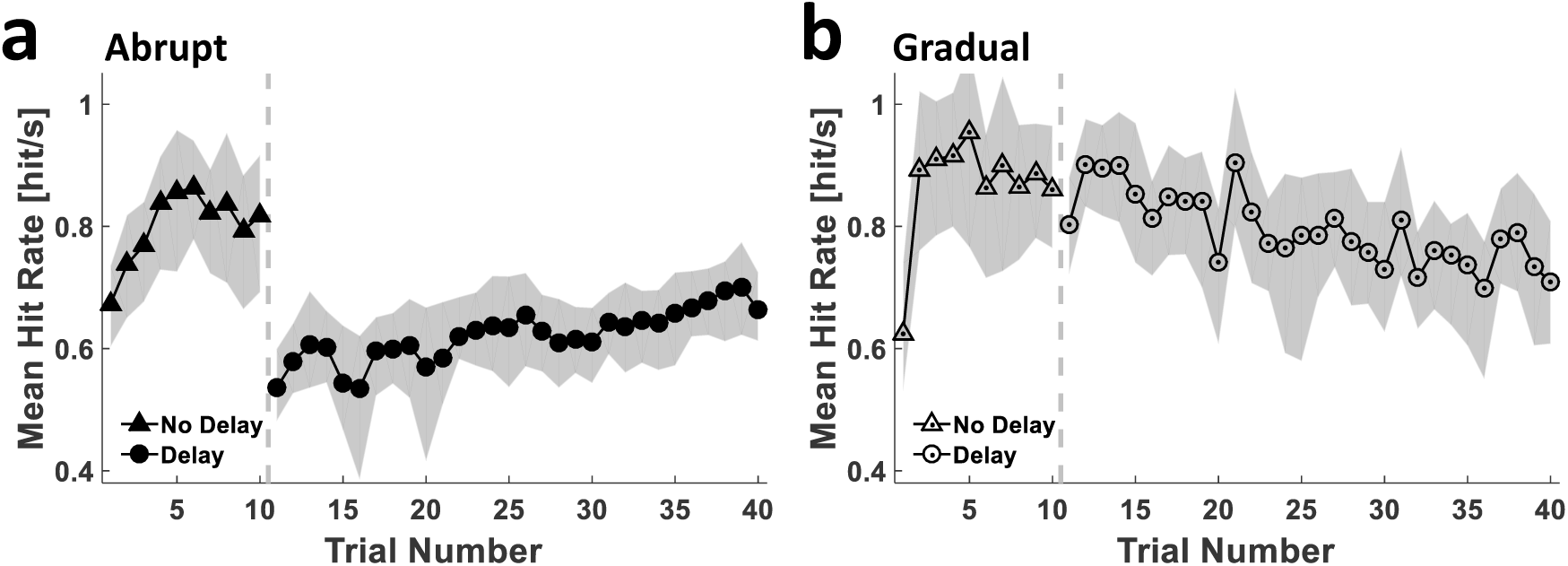
Experiment 2: paddle-ball hit rate in the presence of abruptly-and gradually-introduced delayed feedback.

Time courses of the mean hit rate for all participants in each group of the Abrupt (a, filled markers, N=10) and Gradual (b, hollow-dotted markers, N=10) groups. The grey dashed vertical line separates the **Pong No Delay** (triangles) and the **Pong Delay** (circles) sessions. Shading represents the 95% confidence interval.

Similar to Experiment 1, we examined transfer for each type of schedule of delay presentation to a blind reaching task after each of the Pong sessions (Fig. 2b, blue and orange frames). Analysis of participants’ performance in the blind reaching task revealed that regardless of whether the delay was presented abruptly or gradually, participants made larger reaching movements following the delayed pong game, and the effect size was similar between the two groups. Figure 6a presents the reaching endpoints during the **Post No Delay** and **Post Delay** blind reaching sessions of a representative participant from each group. In both participants, whereas the **Post No Delay** movement endpoints reached close to the targets, the **Post Delay** movement endpoints overshot them. We analyzed the changes in reaching amplitude due to the delayed pong and compared the Abrupt and Gradual groups (Fig. 6b). Playing the delayed pong resulted in a significant increase in reaching amplitudes (main effect of the Session: *F*_(1,18)_ = 19.805, *p* < 0.001, [*mean difference, 95% CI*], 1.638 *cm*, [0.907 2.369]), and this effect was not different between the groups (Session-Group interaction effect: *F*_(1,18)_ = 1.507, *p* = 0.235) (Fig. 6b). These results suggest that the hypermetric blind reaching movements following the experience with the delayed pong were not influenced by the schedule of the delay presentation.

**Fig 6.**
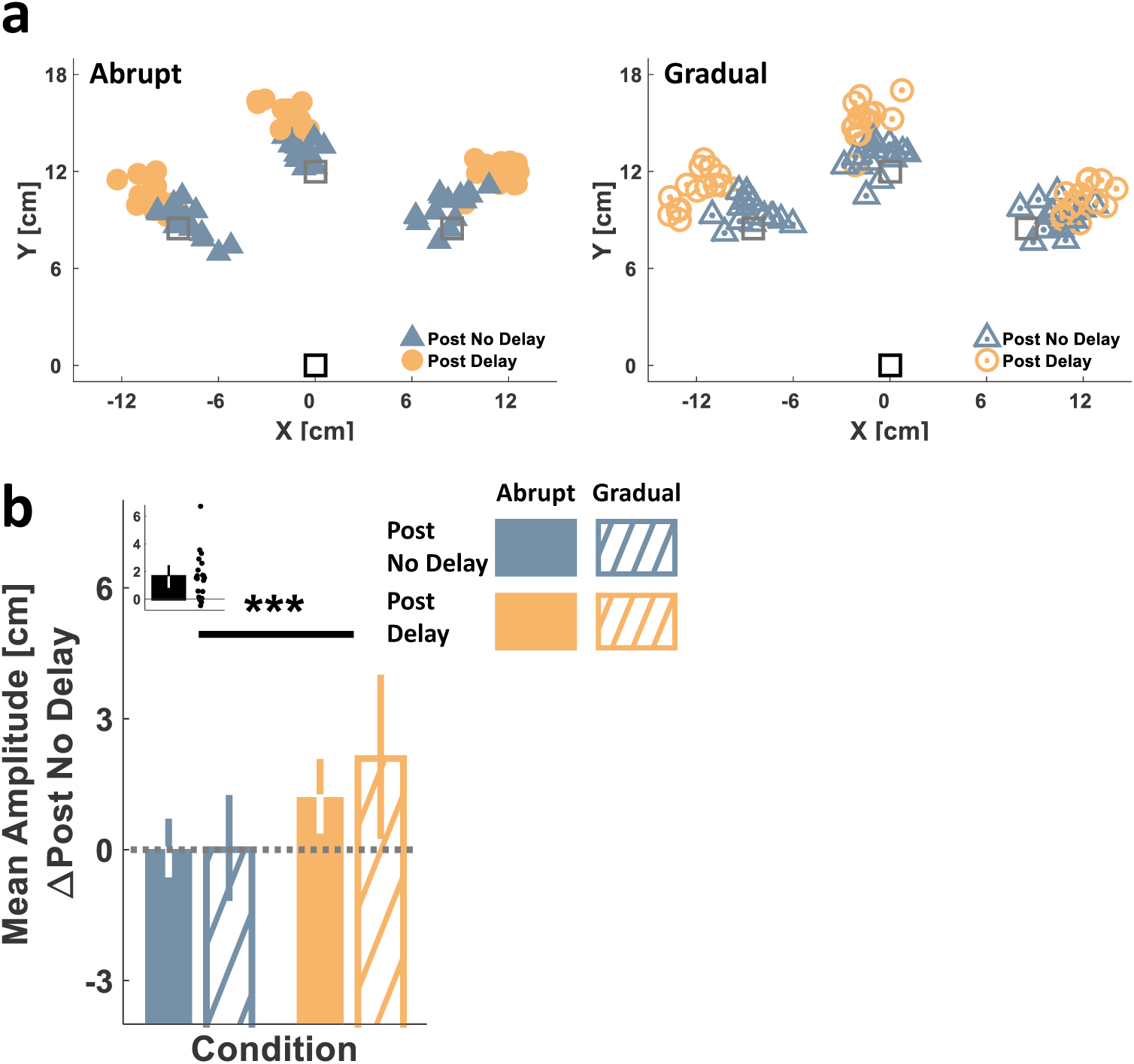
Experiment 2: a comparison between the reaching results in the Abrupt and Gradual groups suggests that the schedule of delay presentation does not influence the representation of delay. (a) Single participant’s experimental results from each of the Abrupt (left, filled markers) and Gradual (right, hollow-dotted markers) groups. Movement start location is indicated by the black square and target locations are marked by the gray squares. Markers represent the end point locations of the hand at movement terminations during the **Post No Delay** (blue triangles) and **Post Delay** (orange circles) **Blind Reach** sessions. (b) Experimental results group analysis. Colored bars represent the mean reaching movement amplitudes towards all targets of each participant, and for each of the **Blind Reach** sessions, averaged over all the participants in each group (Abrupt: filled, N=10, Gradual: diagonal lines, N=10) and following subtraction of each group’s average baseline amplitude. The black bar (inset) represents the difference in mean amplitude between the **Post Delay** and the **Post No Delay** blind reaching sessions for each participant, averaged over all targets and all the participants in both groups. Error bars represent the 95% confidence interval. ***p<0.001.

There was no significant difference in reaching amplitudes between the targets (main effect of Target: *F*(1.327,23.887) = 3.228, *p*_*ε*_ = 0.075). In addition, there was no difference in the change in reaching amplitudes throughout the experiment between the targets (Target-Session interaction effect: *F*(1.517,27.313) = 1.205, *p*_*ε*_ = 0.304), and no difference between the Abrupt and Gradual groups (Group-Target-Session: *F*(1.517,27.313) = 0.114, *p*_*ε*_ = 0.729). Thus, the increase in the blind reaching amplitudes following the delayed pong game was similar across the different targets.

### Experiment 3

#### Transfer of hypermetria to a blind tracking task suggests State Representation as either a Gain or a Mechanical System equivalent rather than a Spatial Shift

Although the comparison between the blind reaching experimental and simulation results suggested that *State* and not *Time* variables were used to represent the delayed feedback, the blind reaching task has two limitations: (1) the increase in blind reaching amplitude following the experience with the delay indicated that the delay affected the representation of the state of the hand, but it may also have masked some extent of the time representation. Since the reaching task is mainly spatial, if there is a partial representation of the time lag it cannot be identified on this transfer task. (2) The blind reaching cannot differentiate between the different types of *State Representations*. All three *State Representation* models – the *Spatial Shift*, the *Gain* and the *Mechanical System* – predict reaching overshoot after the delayed Pong.

Thus, to determine which model best explains delay representation, we conducted an additional experiment in which we examined transfer to blind tracking after each of the **Pong** sessions (Fig. 2c, purple and green frames). On each trial, a target moved along a figure-of-eight path and participants were required to track and maintain the imagined cursor within the target. Importantly, we designed the tracking task so that it would be predictive, and therefore could reveal any temporal components in the representation (for both the *Time* and *Mechanical System Representation* models) (Rohde et al., 2014). To test whether the transfer is influenced by the schedule of delay presentation, the participants were again assigned to one of two groups: Gradual (N=10) and Abrupt (N=10), which were different from the schedule of delay presentation during the **Pong Delay** session.

Predicted tracking performance for each representation model: *Time Representation* (left), *State Representation* (right) – *Spatial Shift, Gain* and *Mechanical System*. The upper panel depicts schematic illustrations of a sinusoidal target trajectory (bold black) and hand trajectories during a tracking task following a non-delayed (**Post No Delay**, dashed gray) and a delayed (**Post Delay**, dotted gray) Pong game. The lower panel depicts the target-hand position space plots for the post non-delayed (**Post No Delay**, purple) and post delayed (**Post Delay**, green) conditions; each corresponds to the target and hand trajectories presented above it. For the *Time Representation* of the delay, the hand trajectory is predicted to precede the target trajectory, resulting in a wider ellipse in the target-hand position space. For the *State Representation* – *Spatial Shift* model, the hand trajectory is predicted to be shifted away with respect to the target trajectory, resulting in an upward shift in the major axis (dashed-dotted dark lines) of the target-hand position space ellipse. For the *State Representation* – *Gain* model, the hand trajectory is predicted to increase in its amplitude with respect to the target trajectory, resulting in an ellipse that has a major axis tilted such that its slope is greater than the slope of the major axis of the **Post No Delay** target-hand position space ellipse. For the *State Representation* – *Mechanical System* model, the hand trajectory is predicted to precede the target trajectory while increasing in its amplitude, bringing about an ellipse that has a major axis tilted such that its slope is greater than the slope of the major axis of the **Post No Delay** target-hand position space ellipse.

Figure 7 presents the predicted blind tracking performance in a single dimension (e.g. sagittal) and for a complete single cycle of the figure-of-eight path during **Post No Delay** and **Post Delay** sessions for each of the representation models. The figure displays both the predicted target and hand position trajectories (upper panels) and the corresponding target-hand position space plots (lower panels). The latter panels depict the position of the hand as a function of the position of the target for each sample during the movement. We assume that during the **Post No Delay** session, the hand lagged slightly behind the movement of the target (Rohde et al., 2014). This relationship is equivalent to an ellipse in the target-hand position space that has a major axis with a zero intercept and a slope of 1. For the **Post Delay** session, if participants coped with the delay using the *Time Representation*, the movement of their hand would be shifted in time with respect to the movement of the paddle, by preceding the path according to the represented time lag. When viewed in terms of the relationship between hand and target, this would result in a wider ellipse in the target-hand position space, and the major axis of this ellipse was expected to overlap with the **Post No Delay** target-hand position space line. Alternatively, if participants represented the delay as a *Spatial Shift*, the entire path of the hand would be shifted farther away from the body of the participant relative to the target. This would result in an upward shift of the target-hand position space ellipse, and thus, a higher ordinate intercept value for its major axis with respect to that of the **Post No Delay** target-hand position space ellipse, but without any change in its slope. A representation of the delay as either a *Gain* or a *Mechanical System* would result in an increase in the hand amplitude from the **Post No Delay** to the **Post Delay** tracking session, which is equivalent to an increase in the slope of the target-hand position space line. Note that a representation of the delay as a *Mechanical System* would also result in a hand trajectory that would precede the target trajectory. Since both the hand lead and hand lag scenarios predict an increase in the width of the target-hand position space ellipse, we examined this temporal relationship using a cross-correlation analysis between the hand and target trajectories rather than based on ellipse fitting.

We analyzed participants’ blind tracking performance by examining the hand and target positional trajectories in both the frontal and sagittal dimensions of the movements. We evaluated the dynamics between hand and target movements by mapping the hand position to the target position for each sample, and by fitting an ellipse to the scatter of each trial. Figure 8a presents examples of target-hand position space scatters and their corresponding fitted ellipses of a single participant from two blind tracking trials – one from a **Post No Delay** session (purple) and one from a **Post Delay** session (green) – and for a single cycle in the sagittal dimension. The results demonstrate that the major axis of the **Post Delay** ellipse had a greater slope than that of the **Post No Delay** ellipse. This type of change in slope is consistent with both the *Gain* and *Mechanical System* representation models, but not with the *Time* or *Spatial Shift* representation models.

For a quantitative analysis of the dynamics between the hand and the target in each of the frontal (Fig. 8b) and sagittal (Fig. 8c) dimensions, we extracted three measures from each trial: the delay between the target and the hand (Target-Hand Delay), the intercept of the major axis of the ellipse (Intercept) and the slope of the major axis (Slope). The Target-Hand Delay was evaluated by finding the lag for which the cross-correlation between the movements of the target and the hand was maximal. Positive values of Target-Hand Delay indicate that the hand movement preceded the movement of the target. The delayed pong did not cause participants to precede their hand movement with respect to the target movement in the blind tracking task. In both the frontal and sagittal dimensions of the task, the mean Target-Hand Delay was not significantly different between the **Post No Delay** and the **Post Delay** blind tracking sessions (Table 1, Session main) ([*mean difference, 95% CI*], frontal: − 0.005, [− 0.021], sagittal: 0.013, [− 0.028 0.002]) (Fig. 8b,c, left). This suggests that participants did not use a *Time Representation* of the experienced delay. Furthermore, participants’ hands did not move farther away from the target in a consistent manner as a result of the experience of the delayed pong. There was no significant difference in the mean Intercept between the **Post No Delay** and the **Post Delay** sessions (Table 1, Session main) (frontal: − 0.541, [−1.093], sagittal: 0.608, [− 0.070 1.286]) (Fig. 8b,c, middle). This suggests that it is unlikely the *State Representation* – *Spatial Shift* model can account for participants’ performance. In contrast, playing the delayed pong game caused participants to execute longer hand movements during the blind tracking task. We found a significantly higher Slope during the **Post Delay** than during the **Post No Delay** session (Table 1, Session main) (frontal: 0.114, [0.046 0.182], sagittal: 0.162, [0.061 0.262]) (Fig. 8b,c, right). These results are consistent with both the *State Representation* – *Gain* and *State Representation – Mechanical System* models.

We did not find an overall difference between the groups in any of these three measures (Table 1, Group main), and no difference in the influence of the delayed pong between the groups (Table 1, Session-Group interaction). These results suggest that similar to the transfer to reaching case (Experiment 2), the schedule of the delay presentation did not influence tracking performance.

### Experiment 4

#### Transfer of hypermetria to a blind tracking task with different movement frequencies suggests State Representation as a Gain rather than a Mechanical System equivalent

The results of Experiment 3 could not dissociate between the *Gain* and the *Mechanical System* representation models. Both models could explain the increase in the movement amplitude during reaching and tracking. However, these models provide different predictions in terms of frequency and velocity dependency.

To illustrate the predicted effect of frequency on movement amplitude, we simulated the frequency response according to each of these representation models (Fig. 9). Since the *Time* and *Spatial Shift* representation models were not associated with any change in the hand amplitude, we focused our simulations solely on the *State Representation* models of the *Gain* and *Mechanical System*. Consider a task where following each **Pong No Delay** and **Pong Delay** session, the hand blindly tracks a target moving along a sinusoid trajectory that has a specific amplitude of 2 cm, but which varies in its frequency. In the case of an accurate tracking performance in the **Post No Delay** session, the hand amplitude would be the same as the target amplitude at all movement frequencies (Fig. 9a, upper panel, magenta lines). For the **Post Delay** session, whereas hypermetria due to a *Gain* representation does not depend on movement frequency, hypermetria resulting from a *Mechanical System* representation is predicted to increase with frequency (Fig. 9a, upper panel, cyan lines). Another way of thinking about this prediction is considering both representations at different velocities. While the *Gain* representation is not expected to depend on movement velocity, the *Mechanical System* representation should yield a velocity-dependent response. Higher frequencies for similar amplitudes of target motion should also result in faster movements of the target.

To test these predictions, in Experiment 4 we examined transfer to a blind tracking task in which the target was moving along a trajectory that was generated as a mixture of five sinusoids (Miall, 1996; Miall and Jackson, 2006), all having the same amplitude (2 cm) but each with a different frequency (0.23, 0.31, 0.42, 0.54 and 0.67 Hz) and with a different phase shift (Fig. 2d, magenta and cyan frames). This served to examine the effect of frequency on the delayed-induced hypermetria.

Predicted effects of tracking movement frequency on the increase in movement amplitude following the delayed pong game. In each of the a-d subfigures, the predictions are presented for the *State Representation - Gain* (left) and *State Representation* – *Mechanical System* (right) models. Upper panels display the **Post No Delay** (magenta) and the **Post Delay** (cyan) amplitudes in cm (a, b) or in dB (c, d), and lower panels present the difference between them. (a, c) When assuming accurate tracking of a target movement that has an amplitude of a 2 cm during the **Post No Delay** session, the *Gain* representation should predict the same increase in movement amplitude for all frequencies during the **Post Delay** session, whereas the *Mechanical System* representation predicts a higher hypermetria with increasing frequency. (b, d) A simulation of an increase in the baseline (**Post No Delay**) movement amplitude with an increase in the movement frequency illustrates that the predictions of both models are equivalent to the predictions for accurate baseline performance when examined in a logarithmic amplitude scale.

We analyzed participants’ blind tracking performance by examining the hand amplitude for each of the main frequencies in the tracking movements and compared the **Post No Delay** to the **Post Delay** sessions. Figures 10a and 10b present the tracking movements of the hand of a representative participant from each session, and the frequency responses, respectively. These figures show an overall increase in the movement amplitude between the **Post No Delay** and the **Post Delay** session and in all five main frequencies. Note that this participant exhibited an increase in the baseline (**Post No Delay**) movement amplitude with an increase in the movement frequency. This effect was also observed in other participants, and in a previous study that examined tracking of a target that moved in frequencies below 1 Hz (Foulkes and Miall, 2000). Because of this effect, the *Gain* model predicts a non-constant increase in the metric measure of the amplitude due to the delay (Fig. 9b). While the predictions between models remain different, this makes them statistically and qualitatively less distinguishable. Therefore, we calculated the 10 ⋅ log _10_ of the resulting power (decibel amplitude, dB) for each movement (Fig. 9c, d, upper panels) and examined the difference between the **Post Delay** and the **Post No Delay** sessions to control for the baseline modulation in the amplitude that results from the increase in movement frequency (Fig. 9c, d, lower panels).

Playing the delayed pong game caused an increase in tracking amplitude, where the magnitude of the increase did not depend on the movement frequency. An analysis of the tracking performance of all participants revealed a significant increase in movement amplitude from the **Post No Delay** to the **Post Delay** session (main effect of Session: *F*_(1,19)_ = 9.423, *p* = 0.006,[*mean difference, 95% CI*], 1.080 *cm*, [0.344 1.816]) (Fig. 10c). Consistent with the results of Experiments 1-3, this effect suggests that the participants did not use either a *Time* or a *Spatial Shift* representation of the delay. As was mentioned above, we also found a significant effect of frequency on the tracking amplitude in both the **Post No Delay** and the **Post Delay** sessions (main effect of Frequency: *F*(2.202,41.830) = 48.199, *p* < 0.001). However, we did not find a dependency of the delay-induced hypermetria on movement frequency (Session-Frequency interaction effect: *F*(2.423,46.040) = 0.132, *p* = 0.910) (Fig. 10d). These results are consistent with a representation of the delay as a *Gain*, rather than as a *Mechanical System* equivalent.

We also calculated the Target-Hand Delay measure using a cross-correlation analysis between the hand and target trajectories for each participant and during each of the **Post No Delay** and **Post Delay** sessions. The Target-Hand Delay is positive when the hand precedes the target. If participants had used *Time Representation* of the delayed feedback, their hand would have preceded the target movement to a greater extent during the **Post Delay** tracking session than during the **Post No Delay** session, and the Target-Hand Delay would have increased, regardless of its baseline level. To a lesser extent, a small increase in this measure would also be predicted by the *Mechanical System* representation (Fig. 7a). We found a significant decrease in the Target-Hand Delay from the **Post No Delay** to the **Post Delay** session (*t*_(19)_ = 3.268, *p* = 0.004). This result indicates that following the **Post Delay** tracking session, participants hand lagged farther behind the movement of the target with respect to the **Post No Delay** session, contrary to the predictions of the *Time Representation* and the *Mechanical System* models.

### *State Representation* of delay as a *Gain* rather than a *Mechanical System* equivalent

(a, b) Single participant’s results. Hand tracking trajectories of a representative participant during the **Post No Delay** (magenta) and **Post Delay** (cyan) sessions (a), and the frequency responses (b). The filled circles represent the amplitude of each of the five main frequencies in the hand trajectories. (c, d) Group analysis. Mean decibel amplitude of all participants (N=20) for each of the five main frequencies (c). The black bar (inset) represents the mean difference in decibel amplitude between the **Post Delay** and the **Post No Delay** blind tracking sessions, and d represents the mean difference separately for each frequency. **p<0.01.

#### Hypermetria during the delayed pong game is consistent with the Gain Representation model rather than the Time, Spatial Shift and Mechanical System models

The hypermetria observed in all the blind transfer tasks that we examined and the finding that its magnitude does not depend on movement frequency suggest that the nervous system constructs a feedforward representation of the delay as a *Gain*. To examine if the representation is also reflected in the pong game, we analyzed the frequency response of the sagittal position trajectories during the game and compared it to the frequency responses predicted by each of the representation models (Fig. 11). For a representative participant, both the sagittal position trajectories of the hand from the last pong trial of each session (Fig. 11a) and the mean profiles of the frequency responses of the last four trials from each session (Fig. 11b) suggest that the participant increased the movement amplitude from the **No Delay** to the **Delay** session. The frequency responses show that the participant had a preferred frequency range of movement. Therefore, to illustrate the predicted effect of each representation model, we simulated frequency responses according to each of the models using the frequency response profile of the no delay session around this frequency range ([0.5 1.5] Hz) (Fig. 11c). Consistent with the simulations of the transfer tasks, the simulation results of the pong game show that the *Time* and *Spatial Shift* models did not predict a change in the movement amplitude due to the delay; In contrast, the *Gain* model predicted a frequency independent hypermetria; and the *Mechanical System* model was expected to result in hypermetria that increases with movement frequency. Such a response is expected to have a stronger effect on higher frequencies, and it might cause one of the higher frequencies to become dominant. Since the participant exhibited hypermetria that did not seem to increase with higher frequencies, his performance is consistent with the *Gain* representation model rather than all the other models that we tested.

All the transfer tasks that we used posed some constraints, such as movements in specific amplitudes and/or frequencies, which enabled us to examine the effect of the delay by controllably compare the performance between the **Post No Delay** and **Post Delay** sessions. For example, the target movement of the tracking transfer task of Experiment 4 directed participants to move in the same five frequencies in both the **Post No Delay** and **Post Delay** sessions, and thus, we could examine the change in amplitude for each of these frequencies. In contrast, because of the less constrained nature of the pong task, participants were not necessarily moving with the same specific frequencies between the non-delayed and delayed pong sessions. Specifically, we found a significant decrease in the dominant movement frequency, which was defined as the frequency with which the movement had the highest amplitude (*t*_(19)_ = 3.708, *p* = 0.002) (Fig. 11d). This means that the participants moved slower in the presence of delay, possibly to reduce the larger magnitude of the spatial disturbance that results from the online delayed feedback in faster movements. For the dominant movement frequency of each session, the respective movement amplitude (maximum amplitude) significantly increased (*t*_(19)_ = −3.879, *p* = 0.001) (Fig. 11e); this effect is not consistent with both the *Time* and the *Spatial Shift* representation models. Moreover, the *Mechanical System* model predicts hypermetria that increases with movement frequency, and thus, this is the only representation that may result in an increase in the dominant movement frequency from the non-delayed to the delayed pong session. Hence, the findings that the dominant movement frequency decreased due to the delay while increasing its amplitude (Fig. 11f) favor the *Gain* model more than the *Mechanical System* representation of visuomotor delay.

**Table 1.** Statistical analyses of the blind tracking task in Experiment 3. For each of the Target-Hand Delay, the Intercept, and the Slope measures, and for each of the frontal and sagittal dimensions of the tracking path, we fit a two-way mixed effect ANOVA model, with the measure as the dependent variable, one between-participants independent factor (Group: two levels, Abrupt and Gradual), and one within-participants independent factor (Session: two levels, Post No Delay and Post Delay). The reported values for each measure are the F ratio, with the corresponding factor and residuals degrees of freedom in parentheses (left column), and the corresponding p-value (right column).

| Effect | Dimension | Measure |  |  |  |  |  |
| --- | --- | --- | --- | --- | --- | --- | --- |
|  |  | Target-Hand Delay |  | Intercept |  | Slope |  |
| | | $F_{(1,18)}$ | $p$ | $F_{(1,18)}$ | $p$ | $F_{(1,18)}$ | $p$ |
| Session main | Frontal | 0.437 | 0.517 | 3.937 | 0.063 | 10.729 | **0.004** |
|  | Sagittal | 2.919 | 0.105 | 3.195 | 0.091 | 9.924 | **0.006** |
| Group main | Frontal | 0.032 | 0.860 | 0.054 | 0.819 | 2.233 | 0.152 |
|  | Sagittal | 0.693 | 0.416 | 1.152 | 0.297 | 0.487 | 0.494 |
| Session-Group interaction | Frontal | 2.949 | 0.103 | 2.322 | 0.145 | 1.110 | 0.306 |
|  | Sagittal | 0.104 | 0.751 | 1.668 | 0.213 | 1.156 | 0.296 |

## Discussion

We exposed participants to delayed feedback in an ecological task – a pong game. Following prolonged experience with the delay, regardless of whether the delay was introduced gradually or abruptly, their movements became hypermetric during subsequent blind reaching and tracking. Simulations suggest that this hypermetria was an outcome of a delay representation as an altered gain rather than as a temporal lag, a spatial shift or a mechanical system equivalent.

### Delay representation – time-based or state-based?

There is an inherent difficulty in deciphering the representation of delay because it is a temporal perturbation that causes spatial effects. For example, a visuomotor delay was shown to increase driving errors (Cunningham et al., 2001) and the size of drawn letters and shapes (Kalmus et al., 1960; Morikiyo and Matsushima, 1990). The ability to determine the representation of the delay in these ecological tasks is limited due to their complexity. Our experimental setup was not entirely natural – the task scene was in 2D and the manipulated objects were not real – and more natural setups are useful for ecological investigations (Jeannerod et al., 1995; Fourneret and Jeannerod, 1998). Nevertheless, the pong game is more complex and dynamic than the motor tasks that are usually employed to study the sensorimotor system: it is composed of multiple interception movements that start and end at various locations of the workspace, and the movement of the target (the ball) is altered by the paddle hits. To overcome the difficulty of extracting the change in representation from such a task, we examined transfer to simple and well-understood tasks. The hypermetria in the transfer tasks implies that the participants used a state-based representation of the visuomotor delay. This may also help accounting for the limited transfer of adaptation to delay to timing-related tasks (de la Malla et al., 2014).

Conversely, recent studies have reported evidence for a time-based representation of delay. In a tracking task, participants adapted to a visuomotor delay by time-shifting the motor command (Rohde et al., 2014), even in highly redundant tasks (Farshchiansadegh et al., 2015). Contrary to our ecological pong game, the tracking tasks in these studies were highly predictable. In addition, the reported time-shift was observed during adaptation and after perturbation removal with a single task. If our participants represented time, it only partially contributed to the adaptation, and was not transferred to blind reaching and tracking. Similar temporal adjustments were also observed with delayed force feedback (Witney et al., 1999; Levy et al., 2010; Leib et al., 2015; Avraham et al., 2017); such adjustments may be based on the capability of sensory organs that respond to force – such as the Golgi tendon organ (Houk and Simon, 1967) or mechanoreceptors in the skin of the fingers (Zimmerman et al., 2014) – to represent delay as a time lag.

### Adaptation to delay versus a spatial shift

There is an apparent similarity between a visuomotor delay and a spatial shift. In previous studies of reach movements, both displaced and delayed feedback caused overshoots that were reduced following adaptation, and a surprise removal of the perturbations caused undershoots (Smith and Bowen, 1980; Botzer and Karniel, 2013). In those studies, participants were required to stop at stationary targets, whereas in the interception task of the pong game, movement endpoints were not constrained. Importantly, in Smith and Bowen (1980), the transfer to movements in the opposite direction was different: overshoot in displacement, and undershoot in delay. This is consistent with our claim that delay is not represented as a spatial shift.

### Mechanical system representation of delay

A dynamic systems approach to the representation of visuomotor delay, and specifically a spring-damper-mass system, was suggested in previous studies (Sarlegna et al., 2010; Rohde and Ernst, 2016; Leib et al., 2017). Unlike our experiments that integrated blind transfer tasks to capture representational changes in feedforward control, the studies that found evidence for the mechanical system representation did so in contexts that included online visual feedback, which may have influenced the motor response (Botzer and Karniel, 2013; Cluff and Scott, 2013).

In studies of tracking tasks, due to the delay, participants changed their grip force control in accordance with the dynamics of a mechanical system (Leib et al., 2017), but the modulations vanished immediately upon delay removal (Sarlegna et al., 2010). Thus, it is unclear whether these effects were the result of a change in an internal representation of hand-cursor dynamics, or an online effect that could possibly be tied to perceptual illusions. The discrepancy between the grip force evidence and ours can also be explained by other results showing that anticipatory grip force adjus™ent is dissociable from trajectory adaptation (Danion et al., 2013).

In terms of kinematics, delay representation as a mechanical system should result in a frequency dependent increase in hand movement amplitude and a lead of the hand with respect to the target (Rohde and Ernst, 2016). Studies of tracking with visuomotor delay observed an increase in task related movement errors (Tass et al., 1996; Sarlegna et al., 2010; Leib et al., 2017) that were modulated with frequency (Langenberg et al., 1998) and a hand-leading phenomenon (Hefter and Langenberg, 1998; Sarlegna et al., 2010; Leib et al., 2017). Since the errors appeared in the presence of perturbed feedback, they could result from online correction attempts (Botzer and Karniel, 2013), and they highlight the difficulty of the sensorimotor system to interpret the delay as an actual time lag. Alternatively, they could have stemmed from changes in both tracking amplitude and the observed temporal phase shifts. However, these studies did not report changes in the movement amplitude due to the delay. Also, hand-leading was not observed in our blind tracking tasks, suggesting that this anticipatory behavior is not part of the feedforward representation of delay.

Recent studies have reported effects of delayed visual feedback on perception – including increased mass (Honda et al., 2013) or resistance (Takamuku and Gomi, 2015) – that are suggestive of a mechanical system representation. The anecdotal verbal responses of our participants that the paddle is “harder to maneuver”, “sluggish”, or “mechanical” are consistent with this view and with previous reports (Smith, 1972; Vercher and Gauthier, 1992). Our findings that the transfer of delay effects is not consistent with explicit reports may stem from the separate processing of visual information for perception and action (Goodale and Milner, 1992).

We considered a representation of a spring-mass-damper that is computationally derived from a Taylor’s series approximation of the delay, and its representational effect is predicted to depend on frequency. In fact, our results of frequency-independent hypermetria are inconsistent with any mechanical system whose gain depends on frequency within the range that we examined; other classes of mechanical systems could yield frequency-independent response, and these would still be consistent with our gain model. Albeit, changes in hypermetria with frequency could still appear for larger movement frequencies, for longer delays or after longer experience. Future studies should examine these possibilities.

### Representation of delay as an altered gain

Our finding that hypermetria during tracking does not depend on movement frequency is consistent with a delay representation as a gain change in visuomotor mapping. Indeed, gain and delay perturbations have several common features: for both of them, the target and cursor locations at movement onset are unaltered, and the aftereffects are similar (Krakauer et al., 2000; Paz et al., 2005). However, the way the magnitude of the spatial effects depends on the movement is different: the effects of delay depend on velocity, and the effects of gain depend on movement amplitude. Indirect evidence for the relationship between gain and delay comes from interference studies. The interference paradigm shows that both successive (Krakauer et al., 1999; Tong et al., 2002; Caithness et al., 2004) or simultaneous (Tcheang et al., 2007; Sing et al., 2009) presentations of competing tasks disrupt learning and consolidation. Delayed visual feedback disrupts adaptation to visuomotor rotation and displacement (Held et al., 1966; Honda et al., 2012), but gain and rotation were not found to interfere with each other (Prager and Contreras-Vidal, 2003). This comparison suggests that gain and delay are processed and represented separately.

Nevertheless, our study provides direct evidence that gain may be used as a representation of delay. None of the previous studies that linked the reported effects of delayed visual feedback to a mechanical system representation (Sarlegna et al., 2010; Honda et al., 2013; Takamuku and Gomi, 2015; Leib et al., 2017) examined them in the context of different movement frequencies or velocities. Because a mechanical system is essentially a frequency-dependent gain together with a phase shift, evaluating the frequency dependency of the representation is critical for distinguishing between the two representations.

### Similar transfer of adaptation between abrupt and gradual schedules

In both reaching and tracking, the strength of transfer did not depend on whether delay was introduced abruptly or gradually. Other studies have reported no difference in the influence of the schedule of perturbation presentation on motor learning of other types of perturbations, either in healthy (Wang et al., 2011; Joiner et al., 2013; Patrick et al., 2014) or in impaired participants (Gibo et al., 2013; Schlerf et al., 2013). In contrast, abruptly-introduced perturbations were shown to strengthen interlimb transfer (Malfait and Ostry, 2004). Furthermore, gradually-introduced perturbations strengthen aftereffects (Kagerer et al., 1997) and the transfer of adaptation to other contexts (Kluzik et al., 2008; Torres-Oviedo and Bastian, 2012). This was found despite the fact that for the same duration of adaptation and for the same maximum magnitude of the perturbation, participants experienced a smaller integral of the perturbation in the gradual compared to the abrupt protocol. In this sense, by comparing the transfer effects with respect to the overall experienced perturbation, and not with respect to its terminal/maximum value, the influence of the gradual presentation of the perturbation on transfer to another context can be considered stronger than the abrupt presentation.

In any case, differences between abrupt and gradual presentations of perturbations may be attributed to the presence or absence of an awareness of these perturbations (Kluzik et al., 2008). Awareness may affect the assignment of the perturbation to extrinsic rather than intrinsic sources (Berniker and Kording, 2008), and to elicit explicit rather than implicit learning (Mazzoni and Krakauer, 2006; Taylor et al., 2014). It may have been the case here that the delay was assigned to an intrinsic source, and that the adaptation to the delayed feedback was a result of an implicit process. This is likely because the brain naturally deals with intrinsic transmission and processing delays. However, this conjecture should be entertained with caution since we probed the delay representation before and after a prolonged exposure to the delay, and therefore, may have missed differences between the abrupt and gradual groups during adaptation.

### The learning rule for adaptation to the delayed pong

Although we saw an improvement in the hit rate in the groups that experienced an abrupt and constant delay, the effects were not strong. In addition, due to the dynamic nature of the gradual protocol, we did not find adaptation in the gradual groups. However, it is obvious in terms of the change in performance during both transfer tasks that an internal representation was indeed constructed during the participants’ experience with the delayed environment and independently of whether they improved or not in the game. Importantly, the findings that the participants were unable to regain their baseline performance during the delayed pong game are likely a direct consequence of the failure to learn the true dynamics of the environment. Although previous studies that examined adaptation to visuomotor delays showed a slight improvement with prolonged training (Foulkes and Miall, 2000), participants could not return to their baseline performance even after five days of exposure to the delay (Miall and Jackson, 2006). In our pong game, only a full representation of the actual time lag between the hand and the paddle could have led to complete compensation of the perturbation and recovery of baseline performance.

The gain representation of the delay is reflected in the change of the participants’ performance during the game: with repeated exposure to the delayed pong, participants increased the movement amplitude. In addition, they exhibited a decrease in the dominant movement frequency. The latter finding can be explained by the influences of the uncontrolled nature of the pong game and the online visual feedback. The pong task does not constrain the participants to continuously track a target that moves with specific frequencies, but it requires to estimate the future locations of the ball and the paddle at each interception attempt. Therefore, participants likely wait for the feedback for planning their next movement, thus reducing their movement velocity. This effect is consistent with evidence that humans slow down their movements when the feedback is delayed (Ferrell, 1965; Avraham et al., 2017), which effectively weakens the delayed-state dependent perturbation.

We did not deal here with the learning mechanisms involved in adaptation to the delay. Various measures can be used to examine adaptation in our pong game (Sternad, 2006; Faisal and Wolpert, 2009; Reichenthal et al., 2016). Since participants were instructed to hit the ball as many times as possible within the time duration of each trial, and were provided with a feedback according to this performance measure, we reported their hit rate throughout the experiments. These hits can be considered as reward signals that influence future interception attempts in a reinforcement learning mechanism (Izawa and Shadmehr, 2011; Wolpert et al., 2011; Shmuelof et al., 2012; Nikooyan and Ahmed, 2015). If the adaptation is error-based (Thoroughman and Shadmehr, 2000; Donchin et al., 2003; Smith et al., 2006; Herzfeld et al., 2014), the candidate error signals need to be identified; for example, the distance between the hand and the paddle at meaningful events during the game such as ball-paddle hits. Further studies are required to understand how the state-based representation of the delay is constructed.

We assumed that the brain uses an estimation of the current position of the hand and updates it according to the delayed visual feedback; thus, for the gain model, it computes a proportional relationship between the hand and the paddle. Another solution that does not require estimation of current hand state is to update a threshold position (Pilon and Feldman, 2006) – set a desired position of the hand that is farther away; this would increase the emergent muscle torques that would bring the arm to the distant position. Also, delayed feedback tends to decrease stability (Milner and Cloutier, 1993), which in turn may change the impedance control of the arm (Burdet et al., 2001). However, this would not cause hypermetria, and such a process may occur in parallel to the update of the internal model (Franklin et al., 2003).

### Representation of longer delays

The representation of visuomotor delay in the sensorimotor system may depend on the magnitude of the delay. Typically, delays in visuomotor integration processes range from 150 to 250 ms (Miall and Wolpert, 1995; Kawato, 1999; Franklin and Wolpert, 2011), and numerous results suggest that humans can cope with such internal delays through neural structures that predict the sensory outcomes of a motor command (Miall et al., 1998; Miall et al., 2001; Imamizu, 2010). The delays that were applied between the hand and paddle movements in our experiments did not exceed 100 ms. For the mean movement frequency that the participants exhibited in the game (∼1 Hz), this absolute delay magnitude is equivalent to a relative delay of ∼10% of the movement cycle duration, which was considered relatively easy to cope with in visuomotor tasks (Hefter and Langenberg, 1998; Langenberg et al., 1998). Thus, it was possibly small enough for the sensorimotor system to be able to adopt a current state-based approximation of the delay to moderately improve in the task. However, higher delays are likely to result in new coping strategies that suggest a time-based representation (Diedrichsen et al., 2007), such as using a delayed state (Witney et al., 1999). Another solution is the move-and-wait strategy (Sheridan and Ferrell, 1963; Ferrell, 1965) where participants move in a feedforward manner in which they stop to wait for the responsive visual feedback, and after the delayed object that is being controlled starts to move, they execute an additional corrective movement. In fact, in the presence of longer delays (from 300 ms to 3.2 sec), the total task completion time is longer (Sheridan and Ferrell, 1963; Ferrell, 1965). We believe that in the context of our pong game, such high delays would deteriorate performance even further, would break down the causal relationship between the motor command and the visual feedback, and would impede any form of representation.

### Implications

Understanding delay representation in the sensorimotor system can be useful for understanding the motor consequences of delay-associated pathologies like multiple sclerosis (Trapp and Stys, 2009). Also, this study opens a new prospect regarding to the role of temporal information in rehabilitation. Traditionally, rehabilitation tasks focus on spatial accuracy. However, reproducing temporal aspects of sensory feedback may improve rehabilitation and help recovering performance at different phases of movements (planning, preparation and execution). Furthermore, our results may be useful in developing in-home rehabilitation procedures utilizing virtual games and simple devices such as a computer mouse. The use of delayed visual feedback as a perturbation has several advantages: it encourages participants to exhibit longer movements, it has a strong transfer to different contexts, and it seems to be robust to explicit processes that would enable to maintain an improvement outside the clinic (Taub et al., 1999).

Understanding the relationship between temporal and spatial aspects of visuomotor coordination is important for the development of additional technologies, such as remote teleoperation (Nisky et al., 2013), brain-machine interfaces (Wolpaw et al., 2000) and prosthetics. The interaction with such systems should be improved by artificially reproducing the natural sensory consequences of the motor commands (Perruchoud et al., 2016), or by incorporating the necessary training if the latter is impossible. Since such systems include substantial feedback delays due to information transmission or processing, the development process of these technologies can benefit from accounting for the spatial aspects in the representation of these temporal discrepancies.

## Author Contributions

GA, RL, AP, LSS, AK, LS, FI, and IN designed the research; GA, RL and AP performed the research; GA and AP analyzed the data; GA, RL, AP, LSS, LS, FI and IN wrote the paper.

## Acknowledgements

The authors would like to thank Ali Farshchiansadegh and Dr. Felix Huang for their help in constructing the experimental setups, Matan Halevi for assistance in data collection, and to Dr. Tirza Routtenberg for her valuable advice on signal processing.

**Authors report no conflict of interest**

